# Rootletin Fiber Dynamics Integrate Cytoskeletal Programs to Shape Neuroepithelial Architecture

**DOI:** 10.64898/2025.12.09.693208

**Authors:** Axelle Wilmerding, Glòria Casas Gimeno, Paula Espana-Bonilla, Susana Usieto, Murielle Saade

**Affiliations:** Cells and Tissues Dept, Instituto de Biología Molecular de Barcelona, CSIC; Parc Científic de Barcelona, Baldiri i Reixac 20, Barcelona 08028, Spain

## Abstract

Central nervous system (CNS) architecture is established early by the organization and proliferative behavior of neuroepithelial (NE) cells, which form a pseudostratified epithelium during neural tube (NT) formation. In neurogenesis, newborn neurons have to detach and exit the neuroepithelium, in a process that requires the coordinated disassembly of apical junctions and the centrosome-cilia module. Although Rootletin—the structural component of the ciliary rootlet and centriolar linker—is classically viewed as a static mechanical element, its behavior in NE cells has not been described in detail. Here, we uncover a conserved, dynamic form of Rootletin fiber organization that remodels in synchrony with NE cell morphogenesis. We show that, in NE cells in interphase, Rootletin fibers extend from the basal body through the apical process toward the nucleus, and that Rootletin maintains its fibrous conformation throughout mitosis. As NE cells initiate apical constriction, Rootletin fibers retract from the apical process and assemble into an anisotropic rim-like structure that aligns with the apical junctional complex. This remodeling is coordinated with microtubule stabilization in low-tension apical endfeet and occurs prior to Lzts1 expression and increased actomyosin contractility. Forced neuronal delamination via Neurogenin-2 or Lzts1 promotes Rootletin rim formation and Rootletin alignment with the apical endfoot cortex. Finally, we show that Zika virus NS5 protein can aberrantly associate with all conformational states of Rootletin fibers, providing a potential mechanical link between ZikaV infection and premature delamination. Together, our findings identify Rootletin as a dynamically regulated cytoskeletal scaffold that orchestrates apical surface remodeling in NE cells and identify a potential mechanism by which ZikaV disrupts neurodevelopment.

## INTRODUCTION

The development of the central nervous system (CNS), including its precise size and shape, is established early on by the distinctive architecture and proliferative capacity of neuroepithelial (NE) cells, which are organized into a highly structured pseudostratified epithelial sheet during neural tube (NT) formation (Bocanegra-Moreno et al., 2023; Miyata et al., 2014; Saade & Marti, 2025). NE cells, whose apical and basal attachments maintain the integrity of the ventricular zone (VZ) (Hatta & Takeichi, 1986; Long et al., 2016), undergo interkinetic nuclear migration (IKNM) in coordination with cell-cycle progression.

After NE cell division, cells committed to neural differentiation detach from the ventricular surface and migrate laterally to the mantle zone (MZ), a critical step for the subsequent formation of functional neuronal circuitry (Frith et al., 2024). This detachment is not spontaneous; it requires a dynamic shape-morphing process in which the apical endfoot progressively constricts, narrowing to a single vertex process before abscission, thereby preserving the integrity of the neuroepithelium (Kawaue et al., 2014; Kawaue et al., 2019). The mechanisms governing neuronal apical constriction and delamination rely on a highly coordinated interplay between adherens junction (AJ) remodeling and cytoskeletal dynamics (Baek et al., 2018; Camargo Ortega et al., 2019; Kasioulis et al., 2017). A downregulation of AJ components, together with contractile actomyosin activity and microtubule reorganization, generates the forces and directionality required for apical constriction (Kasioulis et al., 2017). These cytoskeletal changes also facilitate centrosome disengagement from the ciliary membrane and its passage through the actin-cytoskeletal tunnel toward the detaching apical foot (Kasioulis et al., 2017). Final abscission then separates the ciliary and apical membranes from the apical foot through a mechanism analogous to cytokinesis (Das & Storey, 2014). AJ remodeling and cytoskeletal forces must be precisely coordinated in space and time during neurogenic apical constriction and delamination, and disruptions in these processes have been shown to be causative factors in a diverse array of neuronal migration disorders known as neuronal ectopias (Francis & Cappello, 2021) sometimes linked with conditions of primary microcephaly (Buchman et al., 2010; Doobin et al., 2016; Saade et al., 2020; Turkyilmaz & Sager, 2022).

Known pathological causes of premature neural delamination that impair NT growth include Zika virus (ZikV) infection, which dramatically increases the incidence of neonatal microcephaly by directly targeting human neural progenitor cells (Yuan et al., 2017). Functional analyses in the developing chick NT provide supportive evidence that ZikV non-structural protein NS5 (Ferrero et al., 2019) is sufficient to induce atypical non-genetic ciliopathy, as well as premature apical constriction and delamination and thus premature neurogenesis (Saade et al., 2020). Proteome screening using a human fetal brain cDNA library, combined with imaging of NE cells apical end feet, revealed that ZikV-NS5 interacts with host proteins known that play key roles in centrosome plasticity and ciliogenesis (Saade et al., 2020). One such host target is the ciliary rootlet coiled-coil (CROCC) gene, which encodes the large coiled-coil Rootletin protein (Mahen, 2021).

Rootletin oligomerizes to form to longitudinally aligned filaments with cross-banded striations, projecting from the proximal end of the basal body to form the ciliary rootlet (Ko et al., 2020; van Hoorn & Carter, 2024; Vlijm et al., 2018; Yang et al., 2002). Rootletin fibers have been described to support sensory cilia maintenance (Chen et al., 2015; Gilliam et al., 2012; Styczynska-Soczka & Jarman, 2015; Yang et al., 2005), withstand mechanical forces imposed on basal bodies of motile cilia (Yasunaga et al., 2022), and maintain centrosome cohesion (Au et al., 2017; Conroy et al., 2012; Theile et al., 2023). Rootletin fibers were often described as inert, static structures primarily resisting mechanical stress but without active or flexible properties (Mahen, 2021; Yang et al., 2002). However, recent cryo-electron tomography data revealed that Rootletin fibers possess flexible protrusions and lateral filament networks, enabling some adaptive deformation and tethering to intracellular membranes (van Hoorn & Carter, 2024). Despite these advances, little is known about the organization of Rootletin fibers in NE cells.

In this study, we use the surface epithelium of the developing chick NT to map the subcellular distribution and remodeling of Rootletin fibers across the cell cycle of NE cells. Beyond the centriolar linker/ciliary rootlet axis, we find that Rootletin fibers extend along the apical process towards the cell nucleus and display anterograde and retrograde motion all over the apicobasal axis. Moreover, we find that the Rootletin fiber is not disassembled throughout mitosis. In NE cells undergoing apical constriction, Rootletin fibers retract from the apical process to occupy the apical endfoot, adopting an anisotropic rim-like configuration along the apical junction, coordinating with tubulin stability before Lzts1 expression and actomyosin activation. We show that forced NE cell delamination via transcriptional (induced by Neurogenin-2) or non-transcriptional (induced by Lzts1) mechanisms increases Rootletin fiber rim formation and contact with the apical endfoot cortex, demonstrating that the remodeling of the Rootletin fiber occurs in coordination with the cytoskeletal machinery that drives apical constriction during delamination. Moreover, we find that ZikV-NS5 hijacks Rootletin fibers in all conformational manners, which may be an additional mechanism by which ZikV promotes premature neuronal delamination. Together, we show that Rootletin fiber dynamic remodeling is a key feature of NE cell apical endfoot constriction, which can be disrupted in the context of ZikaV infection.

## RESULTS

### Dynamic Rootletin fiber organization underlies a conserved vertebrate neuroepithelial mechanism

Human Rootletin consists of 2017 amino acids and contains a globular domain and 4 predicted coiled-coil domains (CCD1-4), forming paired bundles between CCD2 and CCD4, but separating in their CCD1 domains to contribute directly to striation of a fibrous structure **(Fig.1A)** (Ko et al., 2020; van Hoorn & Carter, 2024). To investigate the subcellular localization of Rootletin in individual neuroepithelial (NE) cells, we used the chick embryo neural tube (NT), a well-established model system for uncovering fundamental subcellular processes in vertebrate neural development and disease (Saade et al., 2020; Saade et al., 2017; Saade et al., 2013). We introduced low concentrations of a plasmid encoding for tagged forms of the human Rootletin (GFP-Rootletin, Rootletin-mScarlet or Flag-Rootletin) by electroporation, together with the fluorescent-tagged nuclear marker H2B **(Fig.1A, B)**. Transverse NT sections revealed that Rootletin is enriched at the apical endfoot of NE cells, with some fibers extending across the ventricular zone (VZ) along the apico-basal axis **(Fig.1B)**. In interphase, Rootletin localizes at the apical endfoot in NE cells, consistent with its role at the centriolar linker (Theile et al., 2023) connecting the two FOP+ centrioles (yellow arrows) and with the ciliary rootlet beneath the basal body and the Arl13b+ primary cilium (purple arrows, **Fig. 1C-D**), as reported in other cell types (Gilliam et al., 2012; Yasunaga et al., 2022). *En face* imaging across NE cell cycle (**Fig. 1E, F**) revealed that Rootletin maintains its fiber organization during centriole duplication in S phase, tracked using the Centrin marker, which is enriched at the distal lumen of each centriole (Wilmerding et al., 2023) (**Fig. 1E**). In G2, as centrosomes separate, an extended Rootletin fiber localizes to one or both centrosomes (yellow arrows, **Fig. 1F**) and further remains at spindle poles during mitosis, either symmetrically or asymmetrically (yellow arrows, **Fig. 1G**). Furthermore, in transversal sections of the NT, immunostaining of GM130, a cis-Golgi marker (**Fig. EV1A**), reveals that Rootletin organizes fibers along the apical endfoot, adopting a distribution similar to the Golgi, spanning the apical process and extending toward the nucleus (**Fig. EV1B, C**) (Gonzalez-Gobartt et al., 2021; Herrera et al., 2021). We next monitored Rootletin dynamics by live imaging and revealed that Rootletin fiber-like structures that extended along the apico-basal axis move in a manner that is reminiscent of, and suggests synchrony with, interkinetic nuclear migration (white arrows, **Movie EV2**). Together, these findings show that Rootletin in NE cells is not restricted to the centriolar linker; it forms a fiber structure that extends along the apical process and is not static, as it can acquire distinct organizations in NE apical endfeet. Importantly, electroporated Rootletin recapitulated endogenous localization patterns. Immunocytochemistry on chick spinal cord sections (HH24) using a Rootletin-specific antibody confirmed its enrichment at the centriolar linker (yellow arrows, **Fig. EV3A–B**), punctate labeling at the ciliary rootlet, and distribution throughout the VZ (**Fig. EV3C, D**).

**Figure 1.**
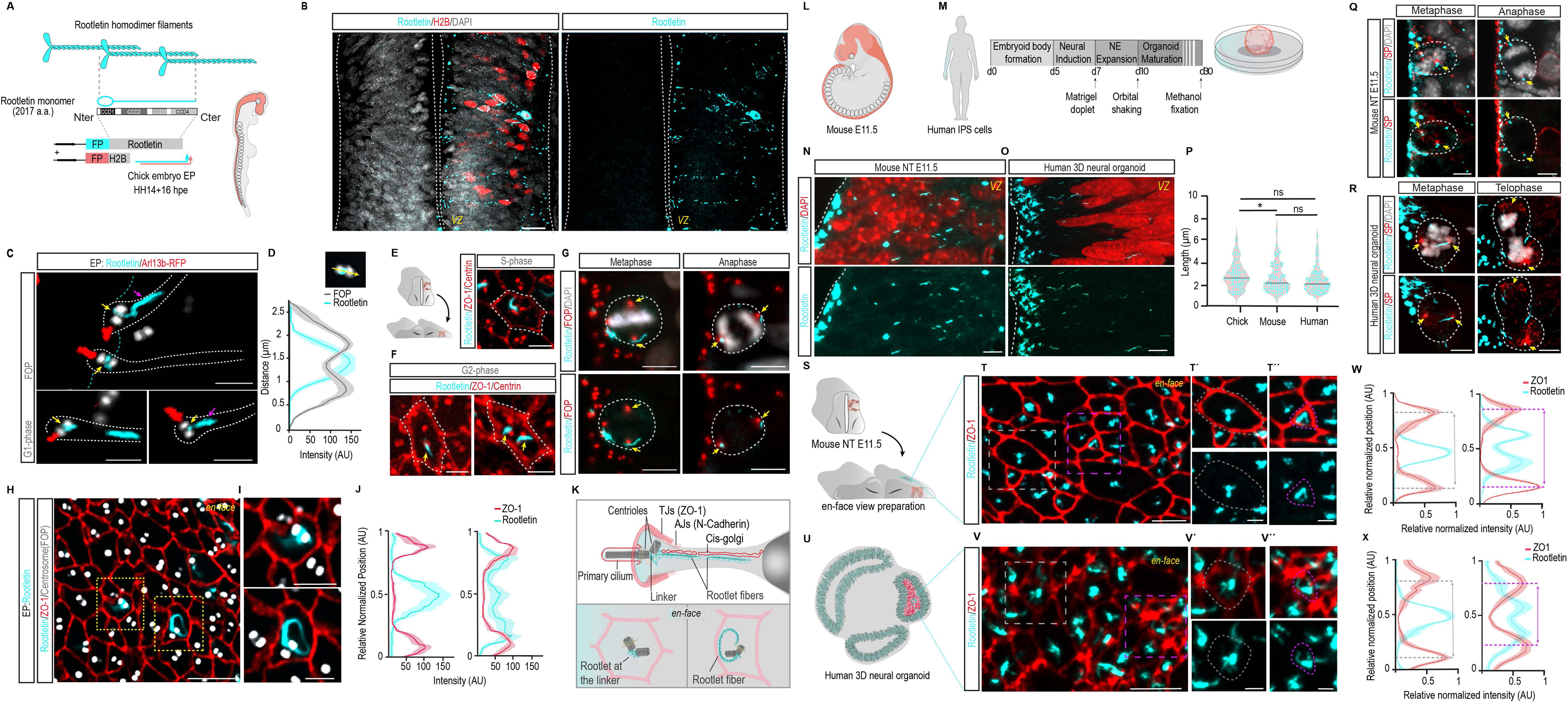
Rootletin forms extended fibers at neuroepithelial cell apical feet and displays conserved subcellular localization across species. **(A)** Schematic depiction of the Rootletin fiber, made of repetitive homodimerized Rootletin. N-terminally tagged Rootletin and the nuclear marker H2B are electroporated into chick neural tubes (NTs) at embryonic stage HH14. NTs are isolated 16 hours post-electroporation (hpe) for imaging analysis. **(B)** Representative images of transversal NT sections show the subcellular distribution of electroporated fluorescent-protein-tagged (FP)-Rootletin and fluorescent-tagged H2B in NE cells at the ventricular zone (VZ), with DAPI-stained nuclei. Dashed lines mark the ventricle and lateral NT edge. **(C)** Representative image of the transversal view of the apical process of electroporated NE cells in interphase, showing EP-Rootletin, Arl13b-RFP-labeled primary cilia, and anti-FOP-stained centrosomes at apical endfeet (white dashed line) along the NT lumen (blue dashed line). **(D)** Fluorescence intensity (AU) profile (AU) of anti-FOP and EP-Rootletin fluorescence intensity across the centrosome of selected NE cells show the location of Rootletin between the two centrioles (n=10). **(E)** Schematic depiction of the chick NT opening from the roof plate for *en face* imaging of NE cell apical endfeet. En face images show the most apical end surfaces of NE cells in S cell cycle phase, with anti-ZO-1-labeled TJs, anti-Centrin-labeled centrioles, and EP-Rootletin subcellular localization. **(F)** Representative *en face* images show the apical endfeet of NE cells in G2 cell cycle phase, with anti-ZO-1-labeled TJs, anti-Centrin-labeled centrioles, and EP-Rootletin subcellular localization (yellow arrow). **(G)** Representative images of mitotic NE cells at metaphase and anaphase, as visualized by DNA staining with DAPI, show EP-Rootletin (yellow arrow) localized at the spindle pole (SP) labelled with anti-FOP. **(H, I)** Representative *en face* images show the apical endfoot of NE cells, showing anti-ZO-1-labeled TJs, anti-FOP-labeled centrosomes, and EP-Rootletin subcellular localization. **(I)** shows insets of **H.** **(J)** Fluorescence intensity (AU) profile plots of anti-ZO-1 and EP-Rootletin across the apical endfoot of selected cells show the two distinct distributions of Rootletin fibers between apical AJs in NE cell endfeet. (n= 10, the apical endfoot length was normalized to 1 and +-SEM is shown). **(K)**The upper scheme shows the apical endfoot organization of an NE cell in interphase, including the polarity complex: adherens junctions (AJs), tight junctions (TJs), the basal body (centrosome) extending the primary cilium into the ventricular lumen, and Rootletin at the centriolar linker and forming a fiber extending toward the nucleus. The two lower schemes illustrate the two observed distributions of Rootletin: on the left, Rootletin localized either at the centriole linker, on the right, forming a fiber that extends toward and aligns with the apical junction. **(L)** Schematic representation of a mouse at embryonic stage E11.5. The central nervous system is highlighted in red. **(M)** Schematic overview of the generation timeline and protocol for day-30 neural organoids derived from a human embryonic stem (hES) cell line. **(N, O)** Representative images of transversal section of mouse NT at E11.5 and human neural organoid at day 30, respectively, showing the subcellular distribution of anti-Rootletin at the VZ with DAPI-stained nuclei. Dashed lines mark the ventricular surface. **(P)** Comparative quantification of Rootletin fiber length at the VZ between the chick embryo NT electroporated with FP-Rootletin, and mouse embryo NT and human neural organoids stained with anti-Rootletin (n=80 fibers for each condition). **(Q, R)** Representative images of mitotic mouse and human NE cells showing immunostained Rootletin (yellow arrow) localized at the SP. Nuclei are DAPI-stained; centrosomes and microtubules are stained with anti-FOP and anti-EB1, respectively. **(S)** Scheme illustrates mouse NT opening from the roof plate for *en face* imaging of NE cell apical endfeet. **(T-T’’)** Representative *en face* image of an E11.5 mouse NT showing anti-ZO-1-labeled TJs, and anti-Rootletin localization. **(T’, T’’).** Enlarged views of the regions highlighted by the gray and purple dashed squares in panel **T**, respectively. **(U)** Scheme illustrates day-30 human neural organoids sectioned for *en face* imaging of NE cell apical endfeet. **(V-V’’)** Representative *en face* view of a human neural organoid showing ZO-1-labeled TJs and endogenous Rootletin localization. (**V’, V’’**) enlarged views of the regions highlighted by the gray and purple dashed squares in panel **V**, respectively. **(W, X)** Fluorescence intensity (AU) profile plots of anti-ZO-1 and anti-Rootletin across the apical endfoot of selected NE cells in mouse NT and human neural organoids, respectively, representative of the two observed distributions of Rootletin. The double-headed arrow indicates the distance between the two peaks of highest anti-ZO-1 intensity. W (left) n=7 cells, W (right) n=7 cells, X (left) n= 7 cells, X (right) n= 7 Cells, the apical endfoot length and fluorescence intensity were normalized to 1 and +-SEM is shown ns: not significant, ^∗^*P* < 0.05. One-way ANOVA. Scale bars 20 μm (**B, C**), 5 μm (**G, H, N, O, Q, R, T, V**), 2 μm (**C, E, F, I, T’’, T’, V’V’’)**

Upon further examination, we noticed that Rootletin at the apical endfoot of electroporated NE cells adopted two apparently distinct organizations. In some apical endfeet, Rootletin is exclusively localized at the centrosome (**Fig. 1H, I (upper), J (left)**). In others, Rootletin appears as a rim-like structure, extending from the centrosome towards, and aligning partially with, the apical junction, where the adherens junctions (AJs), tight junctions (TJs)- and the polarity complex are localised (**Fig. 1 H, I (lower), J (right), K**) (Marthiens & French-Constant, 2009; Saade & Marti, 2025). A similar organization of Rootletin was observed by immunostaining in both cross-sectional (**Fig. EV3E**) and *en face* orientations of the NT, showing a punctuate distribution near TJs at the apical endfoot (**Fig. EV3F, H**). We then performed live tissue imaging using mis-expression of fluorescent-tagged membrane and monitor Rootletin dynamics in an adapted *en face* preparation of cultured chick NT (**Movie EV4**). Tracking Rootletin revealed distinct distributions across apical endfeet: in some, Rootletin localization was restricted to the center of the apical area, where the centrosome is expected to reside (white arrow); in others, emerge fibers in close vicinity of the apical junction (green arrow); and in yet others, a clear rim-like fiber occupying the entirety of the apical endfoot (yellow arrow, **Movie EV4**). Collectively, our data demonstrates that Rootletin in NE cells is not confined to the centriolar linker; it also forms a highly dynamic fiber structure that can extend from the basal body of primary cilia and project upwards along the apical process, or organize in alignment with the apical junction, potentially in a cell-state dependent manner (**Fig. 1K**).

We next examined the subcellular distribution of Rootletin in NE cells of the developing NT across species to assess its evolutionary conservation. For this, we performed immunostaining of the embryonic mouse NT at stage E11.5 (**Fig. 1L**) and human IPSc-derived neural organoids at day 30. (**Fig. 1M**). Transversal sections revealed that Rootletin extends beyond its apical localization to form fibers projecting in both the mouse VZ (**Fig. 1N**) and the human VZ-like (**Fig. 1O**), in a conserved manner that recapitulates the subcellular distribution observed for Rootletin in the chick embryo NT (**Fig. 1P).** In NE cells in mitosis, Rootletin consistently localizes at the spindle pole, either symmetrically or asymmetrically, in both species (yellow arrows, **Fig. 1Q, R**). *En face* imaging of the apical endfoot in mouse (**Fig. 1S, T**) and human (**Fig. 1U, V**) NE cells revealed that the two organizational patterns of Rootletin observed in the chick embryonic NT are conserved; in some apical endfeet, Rootletin was restricted to the centrosome (**Fig. 1T, T’, V, V’**), whereas, in other apical endfeet it appeared as a rim-like fiber aligned with the TJs marker ZO-1 (**Fig. 1T’’, V’’**), as illustrated by fluorescence intensity profiling across the apical endfoot of selected cells (**Fig. 1W, X**). Our data shows that Rootletin exhibits spatially distinct distributions within NE apical endfeet that are conserved across species, underscoring its evolutionary conserved function in NE cell biology.

### Rootletin fiber-mediated rim assembly at the apical membrane coordinates NE cell end-foot constriction

To examine whether the Rootletin fiber undergoes dynamic remodeling in a specific context of the apical endfoot reorganization, we analyzed its subcellular configuration relative to centrosome state and positioning using the marker FOP, and apical endfoot surface using the TJs marker ZO-1, by *en face* imaging 24 hrs hours after HH14 chick embryo NT electroporation (**Fig. 2A-G**). We observed that in some relatively large apical endfeet in which the centrosome is centrally positioned, the Rootletin fiber extends from the centrosome into the cytoplasm (**Fig. 2A**). In other apical endfeet, the Rootletin fiber emanating from the centrosome appears to extend toward and partially curve along the apical junction **(Fig. 2B).** In some others, Rootletin fibers adopt a rim-like configuration that is consistently positioned laterally, toward the periphery of the apical endfoot, in an anisotropic manner (**Fig. 2C**). In apical endfeet where the apical area is apparently reduced and the centrosome is shifted laterally – a hallmark of NE cells undergoing delamination (Saade et al., 2020; Wilsch-Brauninger et al., 2012) - the Rootletin rim aligns completely with the contour of the apical junction (**Fig. 2D, E**). In constricted apical endfeet, in which the centrosome is no longer localized at the most apical end surface and begins migrating basally, another hallmark of NE cells undergoing delamination (Das & Storey, 2014; Kasioulis et al., 2017; Saade et al., 2020), Rootletin signal coalesces and appears concentrated at the apical tip of NE cells (**Fig. 2F**). Quantification of Rootletin fiber length (L2) adjacent to the apical junction relative to the ZO-1 defined apical junction perimeter (L1), a parameter hereafter referred to as Rootlet fiber occupancy of the apical endfoot (AE), revealed that Rootletin fiber occupancy inversely correlates with apical area size (**Fig. 2G**). These observations suggest that Rootletin fiber assembles into an apical rim in concert with NE cell apical endfoot constriction.

**Figure 2.**
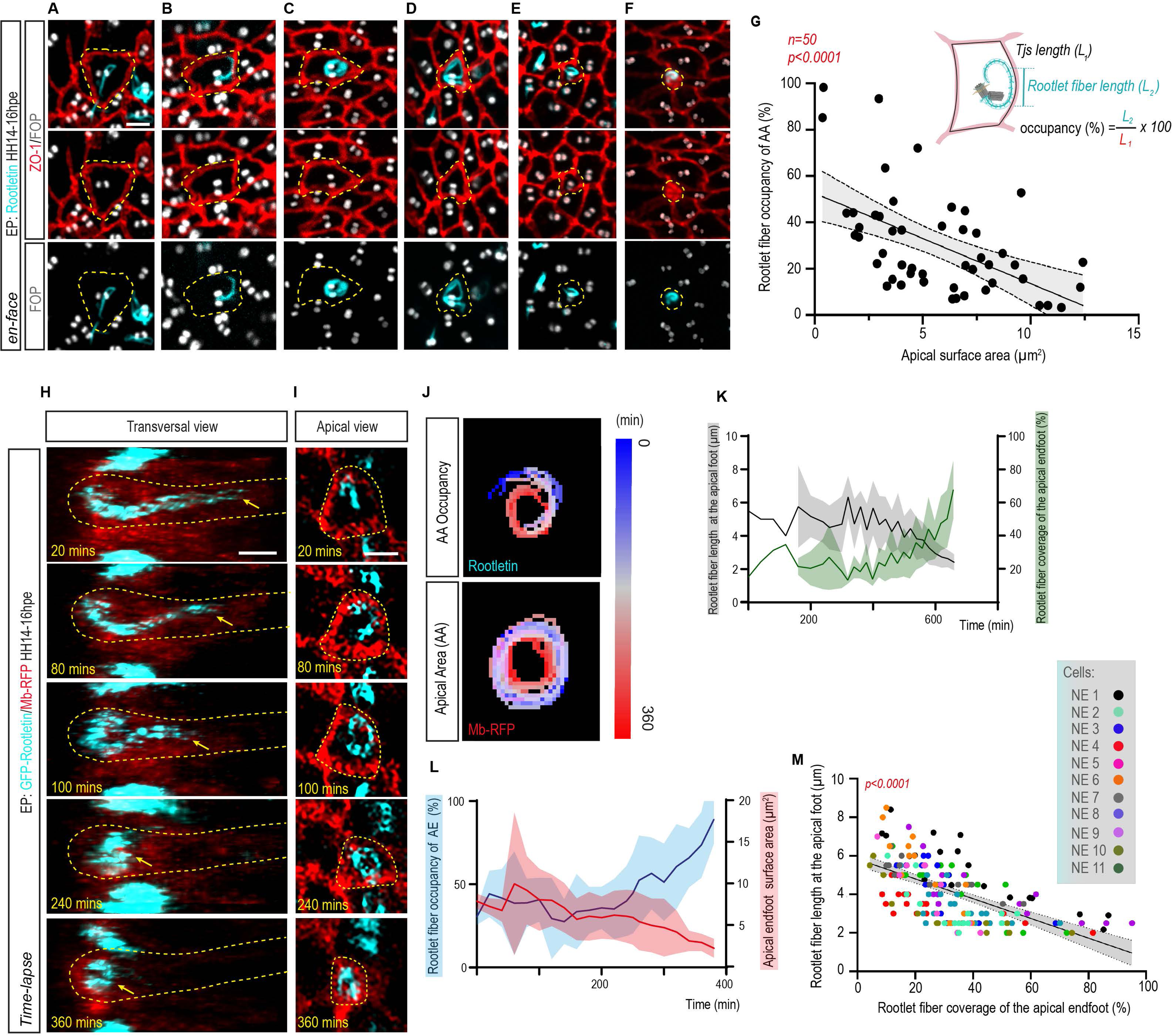
The Rootletin fiber is not static but is actively remodeled into a rim-like structure during NE apical endfeet constriction. **(A-F)** Representative *en face* images showing the localization of the EP-Rootletin fiber in apical *endfeet* of NE cells of the chick embryo NT. TJs are labelled with anti-ZO-1, and centrosomes are labelled with anti-FOP. **(G)** Schematic illustration of an NE cell apical endfoot, with the black line representing the perimeter (L1) of the TJ and the length of the EP-Rootletin fiber (L2) contacting the apical membrane. L1 and L2 parameters were used to measure EP-Rootletin fiber apical endfoot (AE) occupancy. Linear regression analysis revealed a significantly negative correlation between NE cell endfoot apical surface area and EP-Rootletin fiber AE occupancy (P < 0.0001, R² = 0.39). Each point represents one cell (n=50). The regression line is shown with 95% confidence intervals. **(H, I)** Selected frames from a representative *in vivo* time-lapse imaging show a transversal view **(H)** and an *en face* view **(I)** of the apical endfoot of an NE cell co-expressing membrane-RFP (Mb-RFP) and GFP-Rootletin. The yellow arrow indicates the position of the most proximal end of the Rootlet fiber within the apical foot at different time points. Time is indicated in minutes (min), **(Movie EV5)** **(J)** EP-Rootletin fiber AE occupancy and apical endfoot surface area (AA) in (**H, I**) are outlined and temporally color-coded for clarity. **(K)** Measurement of EP-Rootletin fiber length along the apical process (Y axis, gray) of n=11 selected cells shows progressive shortening of the EP-Rootletin fiber over time (X axis). Measurement of the percentage of the apical endfoot area occupied by EP-Rootletin of the same cells (AE coverage,Y axis, green) shows a progressive accumulation of EP-Rootletin at the apical endfoot over time. **(L)** Linear regression shows a significantly negative correlation between EP-Rootletin fiber length along the apical process and percentage of EP-Rootletin coverage of the apical endfoot AE measured in **(K)** (P < 0.0001, R² = 0.38). The values for each of the n=11 cells are color-coded. The regression line is shown with 95% confidence intervals. **(M)** Quantification of apical endfoot surface area (Y axis, red line) over time (X axis) of NE cells undergoing apical constriction (6 of the cells in K, L, plus 5 others, a total of n=11 cells) and quantification of percentage of EP-Rootletin fiber occupancy of the AE occupancy (Y axis, blue line), revealing an increase at the apical end-foot surface over time. \*\*\**P* < 0.0001, F-test (**G, L**). Scale bars 2 μm (**A-E, G, H**).

To further investigate the dynamics of Rootletin fiber remodeling during apical constriction, we monitored GFP-Rootletin and mb-RFP remodeling in electroporated NE cells using live imaging at high spatiotemporal resolution (**Figs. 2H-M**, Movie **EV5**). In 3D reconstructions of transversal z-stacks with 0.5 µm intervals, we observed that the Rootletin fiber progressively retracted over time and became concentrated at the apical endfoot surface (yellow arrow, **Fig. 2H, Movie EV5**), as confirmed by quantification of Rootletin fiber length in 11 apical feet over a 400-minute period (**Figs. 2K, L**). *En face* view of the apical endfoot of the same NE cells revealed a progressive increase in the percentage of the apical endfoot surface occupied by Rootletin, a parameter we termed Rootletin coverage of the AE **(Fig. 2K**). Moreover, we found a strong correlation between these two parameters, indicating that as the Rootletin fiber retracts apically, it is accumulated at the apical endfeet **(Fig. 2L blue line).** As the Rootletin fiber at the apical foot shortened and accumulated at the apical endfoot (**Fig. 2H**) it progressively extended along the apical endfoot membrane (Fig. **2I**, Movie **EV5**). Over ∼100 minutes, the Rootletin fiber progressively adopted a rim-like configuration (Fig. **2I, J**, Movie **EV5**).

We next examined whether the correlation between apical endfoot area and Rootletin AE occupancy we observed in fixed samples (**Fig. 2G**) corresponded to a progressive remodeling of the Rootletin fiber in concert with apical constriction. Indeed, quantification of *en face* time lapses of NE cells undergoing apical constriction (6 of the cells in **Fig. 2K, L**, plus 5 others, a total of 11 cells) revealed that apical endfoot constriction was accompanied by a progressive anisotropic rim-like alignment of the Rootletin fiber with the membrane of the apical endfoot, as measured by Rootletin occupancy of the AE **(Fig. 2J and M, Movie EV5**). Tracking the dynamics of these two structures clearly showed that Rootletin fibers first remodeled into a rim-like structure by ∼100 minutes, after which the apical membrane remodeled accordingly by ∼240 minutes, followed by further constriction of both structures by ∼360 minutes (**Fig. 2I, Movie EV5**). In conclusion, these findings demonstrate that apical constriction of NE cells is accompanied by the dynamic remodeling of Rootletin fibers, suggesting that the progressive transition of the Rootletin fiber into an anisotropic rim-like architecture could play a mechanical role in apical endfoot constriction in NE cells, highlighting Rootletin as a potential key determinant of NE cell morphogenesis.

### Rootletin fiber remodeling coordinates with key cytoskeletal changes during neurogenic apical constriction and delamination

During primary neurogenesis, newborn neurons undergo a marked constriction of their apical endfoot before detaching from the ventricular surface through apical abscission, progressively reducing their epithelial contacts to ensure safe delamination without compromising tissue integrity (Baek et al., 2018; Rousso et al., 2012; Saade & Marti, 2025). This process is driven by dynamic cytoskeletal remodeling and the disassembly of apical junctions (Baek et al., 2018; Das & Storey, 2014; Kasioulis et al., 2017; Rousso et al., 2012). A hallmark of this transition is the upregulation of Lzts1, a microtubule-associated protein which accumulates at AJs in delaminating cells and induces apical constriction downstream of differentiation transcriptional programs (Kawaue et al., 2019, Wilmerding et al., 2021). Consistently, immunostaining of Lzts1 in the *en face* preparations of chick embryonic NT at HH24 (**Fig. 3A**) shows that Lzts1 localizes at the apical junction marked by N-Cadherin, (**Fig. 3A**) and that Lzts1 fluorescence intensity negatively correlates with apical endfoot surface area (**Fig. 3B**), confirming its suitability as a marker of delaminating cells (Kawaue et al., 2019, Wilmerding et al., 2021). We next examined Rootletin fiber organization relative to Lzts1 expression (**Fig. 3C-F**). In Lzts1-apical endfeet, Rootletin fibers adopted an anisotropic rim-like configuration (**Fig. 3C**)—previously shown to be a remodeling step accompanying apical constriction (**Fig. 2I, Movie EV5**). In contrast, neurogenic Lzts1⁺ apical endfeet displayed Rootletin fibers positioned subjacent to the apical junction (**Fig. 3D**), which progressively coalesced as apical constriction advanced and collapsed into a single vertex during neuronal delamination (**Fig.3E**). Quantification showed that Rootletin fiber occupancy of the sub-apical membrane rarely exceeded ∼40% in Lzts1-apices but increased to ∼90% in Lzts1+ apices (**Fig. 3F**). These observations indicate that Rootletin fiber remodeling is tightly associated with neuronal apical constriction and delamination.

**Figure 3.**
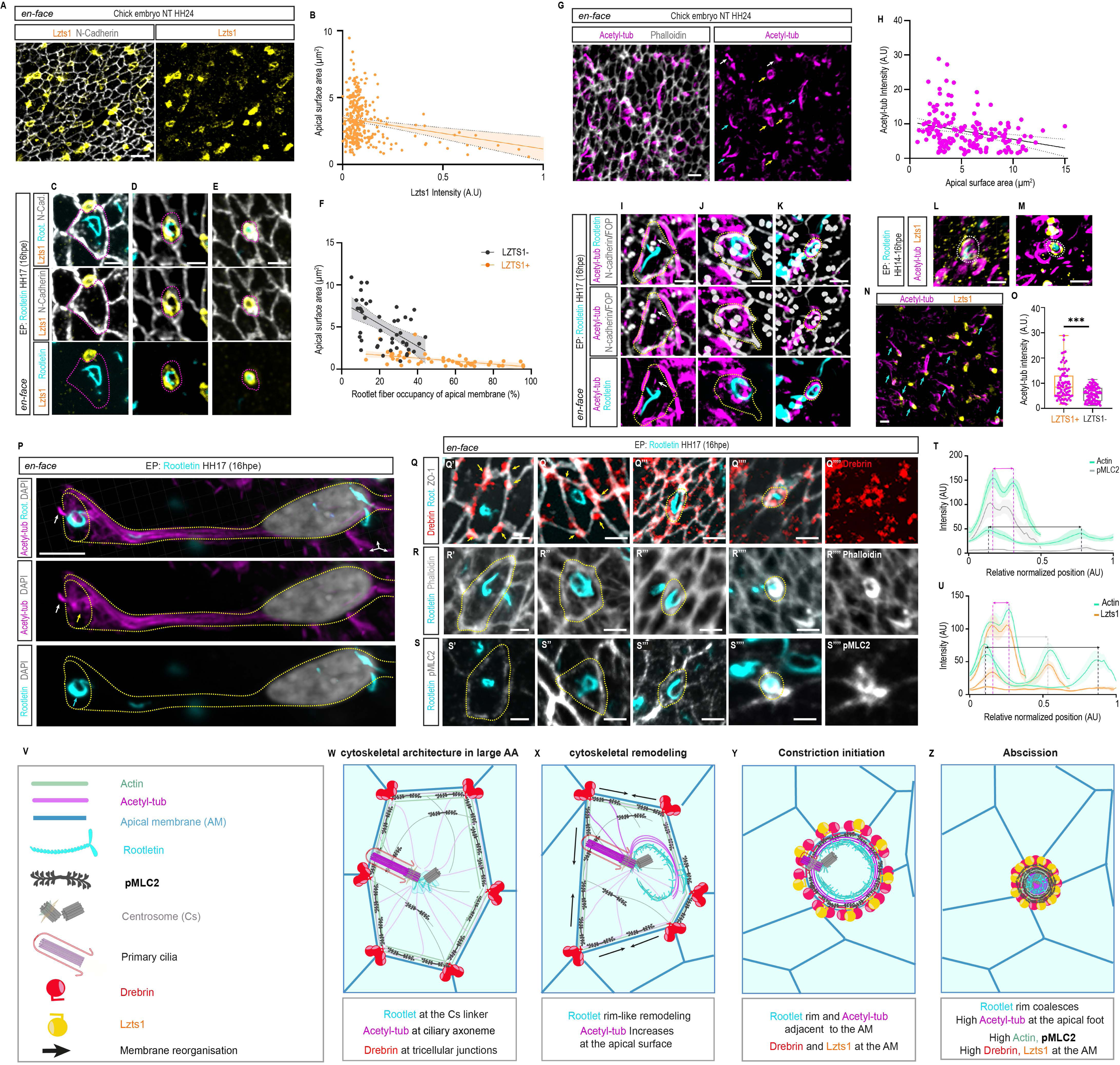
Coordination between Rootletin fiber remodeling and cytoskeletal changes during NE apical endfoot constriction. NE cells require patterned apical cytoskeletal components remodeling to synchronize apical constriction. **(A)** Representative *en face* images of the chick embryo NT at stage HH24, showing the localization of anti-Lzts1 at the apical endfoot cortex in constricted apical endfeet of differentiating cells. NE cells constrict their apical endfoot prior to retraction of their apical process as they delaminate from the VZ. Outline of the apical endfeet is visualized by anti-N-cadherin. **(B)** Scatter plot shows a negative correlation between normalized anti-Lzts1 fluorescence intensity (AU) and the apical endfoot surface area, delineated by anti-N-Cadherin. Each point represents an individual cell. The regression line is shown with 95% confidence intervals. **(C-E)** Representative *en face* images showing the subcellular organization of the EP-Rootletin fiber in apical endfeet of NE cells at HH17 (16hpe), relative to anti-Lzts1 expression. NE cell apical junctions are marked by anti-N-cadherin. Purple dashed lines outline the apical junction of highlighted cells. **(F)** Scatter plot showing the apical endfoot surface area (Y axis) in relation to the EP-Rootletin fiber occupancy of the apical endfoot (AE) (X axis), distinguishing Lzts1− (black) and Lzts1+ (orange) cells. Each point represents an individual cell. The graph shows that Lzts1+ cells have constricted apical endfeet, and that they show stronger correlation between apical endfoot surface area and EP-Rootletin fiber occupancy of the AE. Linear regression lines with 95% confidence intervals (shaded regions) (for Lzts1-, P <0,0001 R²=0,3022, n=45; for Lzts1+, P < 0,0001, R²=0,2642 =, n=48). **(G)** Representative *en face* images of the chick embryo NT at stage HH24, showing anti-acetylated tubulin distribution into short (white arrows) and long (blue arrows) slender structures, as well as rim-like arrangements (yellow arrows). Apical endfeet are delineated by phalloidin staining. **(H)** Scatter plot of anti-acetylated-tubulin fluorescence intensity (Y axis) in relation to apical endfoot surface area (X axis). Each point represents an individual cell. (P <0,0001, R² =0,08738, n=163). The regression line is shown with 95% confidence intervals. **(I-K)** Representative *en face* images showing the subcellular organization of EP-Rootletin fiber in apical endfeet of NE cells at HH17 (16hpe), relative to anti-acetylated-tubulin distribution, centrosome positioning labeled with anti-FOP, and apical junctions labelled with N-Cadherin. The white arrow in **I** indicates primary cilia; yellow dashed lines outline the highlighted apical endfoot. **(L, M)** Representative *en face* images showing the subcellular organization of EP-Rootletin fibers in constricted apical endfeet of neuronally differentiating cells, relative to anti-acetylated tubulin and anti-Lzts1 localization. **(N)** Representative *en face* image of the ventricular surface of the chick embryo NT at HH24 showing the subcellular organization of immunostained acetylated tubulin together with immunostained Lzts1. Cyan arrows highlight acetylated-tubulin enrichment at constricted, Lzts1+ apical endfeet. **(O)** Quantification of anti-acetylated-tubulin signal intensity in Lzts1+ and Lzts1-apical endfeet (n = 163 across 3 embryos; median ± S.D.). **(P)** Representative 3D reconstruction of a transversal view of a NE cell (outlined by a yellow dashed line) at HH17 (16hpe), showing the subcellular organization of EP-Rootletin and immunostained acetylated-tubulin. Cyan and yellow arrows highlight rim EP-Rootlet fibers and enriched Acetylated-tubulin at the apical end surface, respectively. The white arrow marks tubulin acetylation at the primary cilium. **(Q)** Representative *en face* images showing the subcellular organization of EP-Rootletin fibers in NE cell apical endfeet at HH17 (16hpe), relative to anti-Drebrin. Apical endfeet surface areas are delineated by immunostaining against the tight junction (TJ) marker ZO-1. Yellow arrows indicate Drebrin enrichment at tricellular junctions in NE cells with large apical endfeet; yellow dashed lines outline the highlighted apical endfeet. **(R-S)** Representative *en face* images showing the subcellular organization of EP-Rootletin fibers in NE cell apical endfeet at HH17 (16hpe), relative to phalloidin-labelled actin filaments, and anti-Phospho-Myosin Light Chain 2 (pMLC2). Yellow dashed lines outline the highlighted apical endfeet. **(T)** Fluorescence intensity (AU) profile plots of phalloidin-stained actin and anti-pMLC2 across the apical endfoot of selected cells. Double-headed arrows show the separation between apical enrichment peaks of actin and pMLC2 at the apical membrane in large (black) versus small (purple) apical endfeet (n=6 cells; The apical endfoot length was normalized to 1 and +-SEM is shown). **(U)** Fluorescence intensity (AU) profile plots phalloidin-labelled-actin and anti-Lzts1 at the apical endfeet of selected NE cells. Double-headed arrows indicate the distance between enrichment peaks of actin and Lzts1 at the apical membrane in large (black), intermediate (grey) and small (purple) apical endfeet (n= 6 cells; the apical endfoot length was normalized to 1 and +-SEM is shown). **(V)** Key structures, cytoskeletal components and organelles highlighted in (W-Z) (**W-Z**) Model illustrating cytoskeletal rearrangements driving apical constriction during neuronal delamination. First, in low-tension apical endfeet, Rootletin fiber remodeling coordinates with tubulin stability to predefine the future apical endfoot. Then, there is an enrichment of Lzts1 and spread of Drebrin around the apical endfoot cortex, and subsequent increase in contractile actomyosin activity. For (O), Mann–Whitney test, ***P < 0.0001. Scale bars 10 μm (**P**), 5 μm (**A, G, N**), 2 μm (**C-E, I-M, Q-U).**

We next examined how Rootletin fiber remodeling coordinates with other cytoskeletal components-specifically microtubules and the actomyosin network – whose interplay shapes the spatiotemporal dynamics of apical abscission during primary neurogenesis (Kasioulis et al., 2017). We first performed immunostainning of the stable microtubule marker, acetylated α-tubulin in the chick embryonic NT using *en face* imaging. This revealed apically-oriented short slender structures (<10 μm, white arrows), likely corresponding to the axoneme of primary cilia (**Fig. 3G**) (Huangfu & Anderson, 2005), as well as longer basally-oriented bundles (blue arrows) and in some apical endfeet, a sub-apical rim-like organization (yellow arrows) (**Fig. 3G**). Acetylated tubulin intensity at the apical endfoot negatively correlated with apical endfoot area (**Fig. 3H**), suggesting dynamic regulation of microtubule stability during constriction and delamination.

We then analyzed Rootletin fiber patterns and microtubule stability at the apical endfoot by *en face* imaging, and staining centrosomes with anti-FOP and apical junction with N-Cadherin (**Fig. 3I-K**). In large apical endfoot with centrally positioned centrosomes and extended ciliary rootlet fibers, acetylated microtubules were enriched in primary cilia (**Fig. 3I**). In apical endfeet where Rootletin fibers formed an anisotropic rim, acetylated microtubules accumulated along the same surface as the rim (**Fig. 3J, P**). In constricted apical endfeet with laterally displaced centrosomes, acetylated microtubules concentrated subjacent to the apical junction, overlapping with the Rootletin rim (**Fig. 3K**). In Lzts1⁺ cells, both acetylated microtubules and Rootletin rims localized sub-apically (**Fig. 3L**). Upon full constriction of Lzts1⁺ apical endfeet, Rootletin fibers collapsed into a single vertex, and stable α-tubulin became broadly distributed along the slender apical process under delamination (cyan arrows) (**Fig. 3M, N**). Moreover, acetylated microtubule intensity was significantly higher in Lzts1+ compared with Lzts1-apical endfeet (**Fig. 3O**), confirming a substantial reorganization of microtubule stability during neurogenic apical constriction and delamination.

Detachment of newborn neurons from the neuroepithelium depends on the coordinated activity of actin and microtubules, mediated in part by the cross-linking protein Drebrin, which has been proposed as a key regulator of this terminal cytoskeletal network (Kasioulis et al., 2017). In addition, in other cellular contexts, Drebrin has been shown to play a key role in cell-cell junctions ensuring robust intercellular adhesion and tissue organization (Rehm et al., 2013). Hence, we examined Drebrin distribution relative to Rootletin fiber organization (**Fig. 3Q**). In large apical endfeet, where Rootletin fibers were centrally positioned or formed an anisotropic rim, Drebrin accumulated at tri-cellular junctions (yellow arrows), regions of high mechanical tension that require robust cytoskeletal support (Bosveld et al., 2018) (**Fig. 3Q’, Q’’**). In small apical endfeet with sub-apical Rootletin fibers, Drebrin became uniformly distributed along the apical junction (**Fig. 3Q’’’)** and remained so in constricted apical endfeet where Rootletin fibers coalesced (**Fig. 3Q’’’’**).

In NE cell apical endfoot, actomyosin is primarily distributed circumferentially along the apical membrane with a faint medioapical network (Alvarez et al., 2024; Martin & Goldstein, 2014, Ranie & White, 2025). In newborn neurons, circumferential actomyosin networks have been reported to drive apical constriction and delamination (Kasioulis et al., 2017). To characterize actomyosin organization in NE apical endfeet, we stained F-actin with phalloidin and detected active myosin using an antibody against the phosphorylated form of Myosin Light Chain 2 (MLC2) at Thr18 and Ser19 (**Fig. 3R, S**), and measured their intensity across apical endfeet (**Fig. 3T**). In relatively large NE apical endfeet with centrally positioned or rim-localized Rootletin fibers, F-actin and active myosin show a predominantly circumferential disposition, with minimal medioapical signal (**Fig. 3R’, R’’, S’, S’’)**. Constricted NE apices with Rootletin fibers lining the apical cortex present relatively weak F-actin and active myosin signals, (**Fig. 3R’’’, S’’’**) which sharply-increases as apical endfoot shrank and Rootletin fibers coalesce (**Fig. 3R’’’’, S’’’’)**). Combined phalloidin with anti-Lzts1 staining confirmed this pattern (**Fig. 3U**); F-actin levels were low in Lzts1-apices, and increased in concordance with Lzts1 levels in more constricted Lzts1+ apical endfeet (**Fig. 3U**).

Together, these experiments delineate the spatiotemporal choreography of cytoskeletal remodeling during neurogenic apical constriction and delamination (**Fig. 3V, W-Z**). In expanded NE apices, stable microtubules concentrate at the ciliary axoneme, Rootletin fibers extend toward the nucleus, actomyosin contractility is low-potentially maintaining central centrosome positioning (Jimenez et al., 2021) - and Drebrin localizes at tricellular junctions (**Fig. 3V, W**). Upon neurogenic commitment, stable microtubules assemble and Rootletin fibers remodel into rim-like conformation to one corner of the apical endfoot, reflecting an anisotropic organization (**Fig. 3V, X**). As constriction proceeds, the apical endfoot membrane constricts around the microtubule and Rootletin fibers, Lzts1 and Drebrin redistribute along the apical endfoot cortex, actomyosin contractility increases, and the centrosome shifts laterally (**Fig. 3V, Y**). Finally, Rootletin fibers coalesce into a single vertex, as stable microtubules redistribute along the narrowing apical foot before abscission (**Fig. 3V, Z**). This sequential coordination of cytoskeletal remodeling ensures proper dynamics of neurogenic apical constriction and delamination without compromising the tissue integrity of the neuroepithelium.

### Coupling of Transcriptional and Mechanical Programs Drives Rootlet Remodeling Before Delamination

To determine whether Rootletin fiber remodeling precedes neuronal delamination in the embryonic chick NT, we first mis-expressed fluorescent-tagged membrane (pCS2-mb) together with proneural factor Neurogenin 2, whose ectopic expression (pCAGGS-Neurog2-IRES-nucGFP, pCAGGS-Neurog2), induces premature differentiation of NE cells into neurons (**Fig. 4A-L)**. Neurog2 is known to transcriptionally activate RND2 to coordinate apical actomyosin activation, and to indirectly repress cadherins and other AJ-associated molecules (**Fig. 4A**) (Heng et al., 2008; Kawaue et al., 2019; Pacary et al., 2011; Pacary et al., 2012). *En face* imaging and measurements of NE cell apical endfeet revealed significantly smaller apical endfeet surfaces in Neurog2-EP cells compared to controls (**Fig. 4B. C**). To confirm neuronal commitment, mis-expression of Neurog2 was combined with the pSox2-EGFP and pHUC/D-RFP reporter constructs to track NE cell identity (**Fig. 4D, E**) (Saade et al., 2013, Park et al. 2000). As expected, FACS analysis at 24 hpe showed a significant increase in neurons (pHUCD-RFP+) accompanied by a reduction in NE progenitors (pSOX2-GFP+) in Neurog2-EP embryos compared to controls (Fig. **4D, E**).

**Figure 4.**
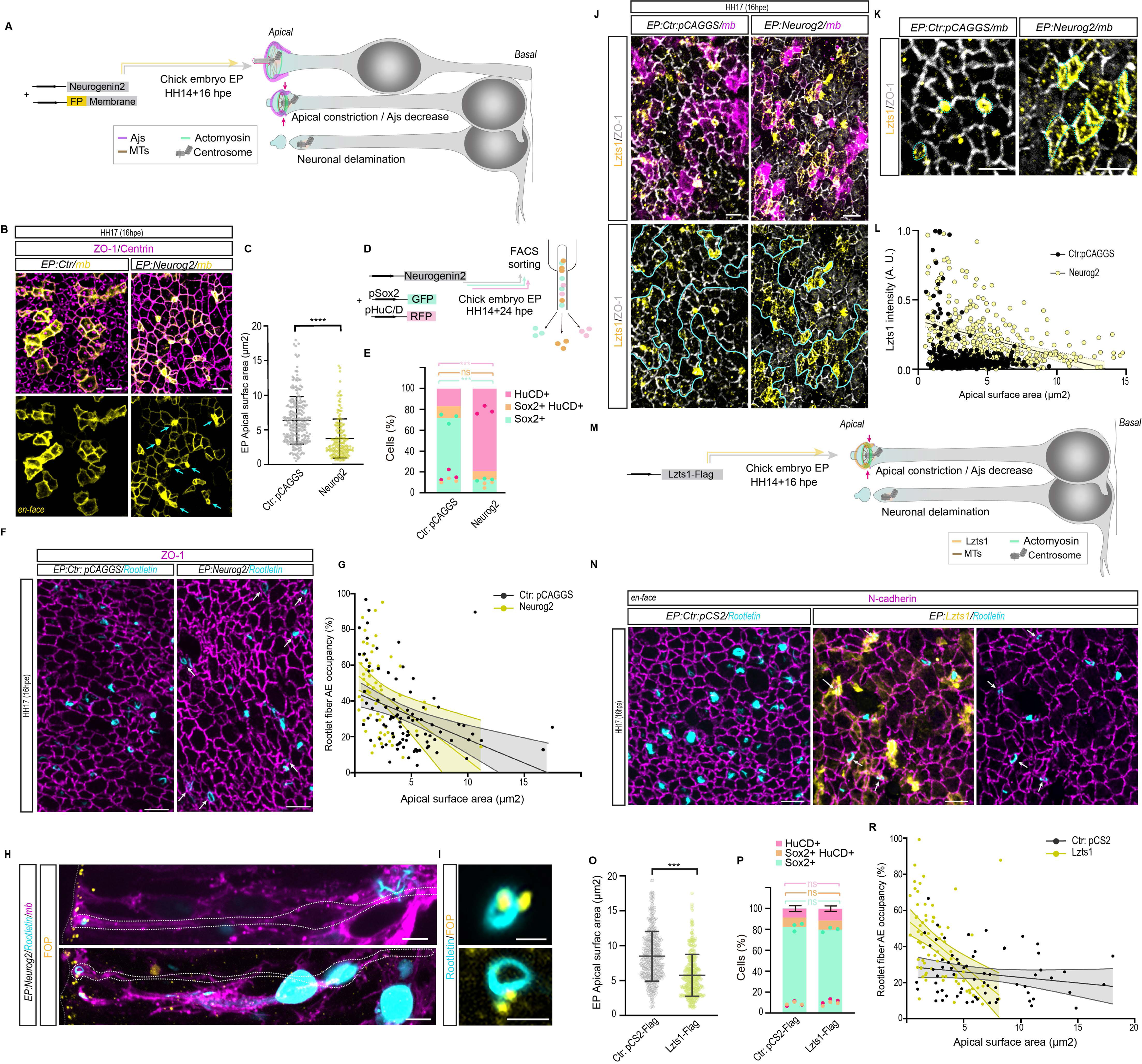
Forcing neuronal delamination actively remodels Rootletin fibers at the apical endfeet. **(A)** Schematic showing the experimental setup: electroporation (EP) of plasmids encoding for the proneuronal factor *Neurog2* (pCAGGS-Neurog2-IRES-nucGFP, or pCAGGS-Neurog2) together with fluorescent tagged pCS2-membrane in the developing chick NT at HH14, for fixing and immunostaining at HH17 (16 hpe). Scheme illustrates NE cell behavior following Neurog2 EP: cells enter G0, undergo sequential apical constriction coordinated by microtubules (MTs) and the actomyosin cable, increase actomyosin contractility, downregulate adherens junction (AJs), move the centrosome basally, detach the apical foot, and delaminate. **(B)** Representative *en face* images of chick embryo NT electroporated with empty vector pCAGGS (control Ctr) or Neurog2, both in combination with pCS2-mb and immunostained for TJ marker ZO-1 and the centriole marker Centrin. Cyan arrows indicate constricted apical endfeet. **(C)** Quantification showing a decrease of the apical endfeet surface area in Neurog2 EP NE cells compared with Ctr pCAGGS EP cells. **(D)** Schematic of experimental set up: co-EP of empty pCAGGS (Ctr) or Neurog2 with the two fluorescent reporters pSOX2-GFP and HUCDp-RFP. Embryos were electroporated at HH14 and harvested at 24 hpe for FACS **(E)** Quantification of the percentage of reporter-expressing cells corresponding to each cell type: NE progenitor (GFP+, green), transitional cells undergoing neurogenic differentiation (GFP+, RFP+, orange), and neurons (RFP+, pink), in embryos electroporated with either pCAGGS (Ctr) or Neurog2. Data represent mean ± SEM (n=4 embryos per replicate, 3 replicates per condition). **(F)** Representative *en face* imaging of NE cell endfeet EP with pCAGGS or Neurog2, in combination with EP-Rootletin and immunostained for ZO-1. **(G)** Scatter plot showing apical endfoot surface area (X axis) as a function of EP-Rootletin fiber occupancy of the AE (Y axis) in Ctr:pCAGGS (black) versus Neurog2 (cyan) EP cells. Each point represents an individual cell. Linear regression lines with 95% confidence intervals (shaded regions) show a negative correlation between EP-Rootletin fiber occupancy and apical endfoot surface area, more pronounced in Neurog2 EP cells. Neurog2 EP cells show uniformly reduced apical end surface areas, with EP-Rootletin fibers occupancy levels extending toward higher percentages. (for Ctr pCAGGS, P < 0.0001, R² = 0.1529, n= 92 cells; for NeuroG2: P = 0.007, R² =0,1205, n= 93 cells from 3 experiments, from n=4 embryos per experiment). **(H, I)**. Representative transversal images of an NE cell (outlined by a white dashed line) mis-expressing Neurog2, fluorescent-tagged mb in combination with Rootletin, showing Rootletin fiber rim associated to the anti-FOP+ centrosome at the subapical domain. **I** insets of **H.** **(J, K)** Representative *en face* images of NE endfeet mis expressing pCAGGS or Neurog2, in combination with fluorescent-tagged mb, stained with anti-ZO-1 and anti-Lzts1. Blue lines in the lower panels delineate EP apical zones defined by fluorescent-tagged mb in upper panels, showing high Lzts1 expression in Neurog2 EP apical endfeet compared to controls. **K** insets of **J** highlighting premature Lzts1 expression in Neurog2 EP cells. **(L)** Scatter plot showing apical endfoot surface area (X axis) as a function of anti-Lzts1 fluorescence intensity (AU) at the apical membrane (Y axis) in Ctr:pCAGGS (grey) versus Neurog2 (light yellow) EP cells. Each point represents an individual cell. Linear regression lines with 95% confidence intervals (shaded regions) show a negative correlation between the two parameters (for pCAGGS P < 0.0001, R² =0,06757, n=422 cells; for NeuroG2: P < 0.0001, R² =0,1776, n=366 cells from 2 experiments, from n=3 embryos per experiment). Note that in Ctr EP cells, Lzts1 expression is restricted to small apical surface areas, whereas Neurog2 EP cells display higher Lzts1 intensity extending to larger apical surface areas. **(M)** Schematic of experimental set-up showing mis expression of pCS2-Lzts1-Flag in the chick NT at HH17 (16hpe). Lzts1 localized at AJs functions by both activating actomyosin contractility and inhibiting MTs assembly at AJs, orchestrating cytoskeletal rearrangements that drive apical constriction and neuronal delamination. **(N)** Representative *en face* images of NE cell endfeet EP with pCS2 or Neurog2, in combination with FP-Rootletin and immunostained for N-cadherin. **(O)** Quantification showing a decrease of the apical endfeet surface area in Lzts1-Flag + Rootlein EP NE cells compared with Ctr pCS2-Flag + Rootletin EP cells. Each point in the graph represents an individual cell; bars represent mean +- SEM. Unpaired t-test P < 0.0001 **(P)** Quantification of the percentage of reporter-expressing cells corresponding to each cell type: NE progenitor (GFP+, green), transitional cells undergoing neurogenic differentiation (GFP+, RFP+, orange), and neurons (RFP+, pink), in embryos electroporated with pCS2-Flag (Ctr) or pCS2-Lzts1-Flag. Data represent mean ± SEM (n=4 embryos per replicate, 3 replicates per condition). **(Q).** Representative en face image of the chick embryo NT electroporated with either pCS2-Flag or pCS2-Lzts1-Flag and immunostained against ZO-1. **(R)** Scatter plot showing apical endfoot surface area (X axis) as a function of EP-Rootletin fiber occupancy of the AE (Y axis) in Ctr:pCS2-FLAG (black) versus pCS2-Lzts1-Flag (yellow) EP cells. Each point represents an individual cell. Linear regression lines with 95% confidence intervals (shaded regions) show a negative correlation between EP-Rootletin fiber occupancy and apical endfoot surface area, more pronounced in Lzts1 EP cells. Lzts1 EP cells show uniformly reduced apical end surface areas, with EP-Rootletin fiber occupancy levels extending toward higher percentages. (For pCS2-Flag P = 0,1358, R² =0,03443, n=71 Cells; for Lzts1-Flag P < 0.0001, R² =0,2268, n=76cells from 3 experiments, from n=4 embryos per experiment). For **C, E, O, P,** ns: not significant, ***P < 0.0001. Scale bars 5 μm (**B, F, H, J, K, N, O**), 2 μm (**I**).

We then examined Rootletin fiber distribution in Neurog2+ NE cells in *en face* preparations of EP chick NT and found that they adopt a rim-like configuration adjacent to ZO-1 in cells with constricted apical endfeet (**Fig. 4F**). Moreover, quantification revealed that Neurog2+ constricted apical endfeet show higher Rootletin fiber occupancy at the apical endfoot (AE) compared to controls (**Fig. 4G**). In transversal sections, Neurog2+ cells displayed the centrosome-Rootletin rim complex positioned in the sub-apical region (**Fig. 4H**), indicating that Rootletin fiber assembly into an apical rim precedes neuronal delamination.

As previously reported, Neurog2 upregulates Lzts1 through proneural downstream networks rather than direct transcriptional regulation (Kawaguchi, 2020; Kawaue et al., 2019). Indeed, when we ectopically introduced Neurog2 by electroporation we observed that, while in *en face* preparations of the control condition, Lzts1 expression was restricted to the most constricted apical endfeet, in Neurog2 mis-expression, EP NE cells showed a marked increase of Lzts1 expression in cells with still expanded apical endfeet, indicating a premature initiation of the delamination program (**Fig. 4J, K, L**).

To further reproduce the premature presence of Lzts1, we mis-expressed Lzts1 (pCS2-Lzts1-Flag) in NE cells of the chick embryo NT to test whether AJ-mediated mechanical execution alone could promote Rootletin fiber remodeling before neuronal delamination (**Fig. 4M**). Lzts1 gain-of-function has been shown to promote apical constriction and neuronal delamination by activating cytoskeletal dynamics and weakening cell-cell adhesion (**Fig. 4M**). Indeed, *en face* analysis confirmed significantly smaller apical endfoot surfaces in Lzts1 electroporated cells compared to controls (Fig. **4N, O**). However, when Lzts1 mis-expression was combined with pSox2-EGFP/pHUCD-RFP reporters, the proportions of NE progenitors (pSOX2-GFP+ cells) and neurons (pHUCD-RFP+ cells) remained unchanged relative to controls (**Fig. 4N, O**). This was expected since Lzts1 induces mechanical apical constriction by targeting the cytoskeleton and weakening AJs, but does not initiate the neurogenic program (Wilmerding et al., 2021).

Finally, we assessed Rootletin fiber organization in Lzts1-EP NE cells and found that, similarly to Neurog2, Lzts1 mis-expression caused Rootletin fibers to adopt a rim-like conformation adjacent to ZO-1 in constricted apical endfeet (**Fig. 4Q**). Quantification confirmed increased Rootletin fiber occupancy of the AE in Lzts1+ constricted apices, preceding delamination, compared with controls (**Fig. 4R**). Together, these results demonstrate that transcriptional commitment (via Neurog2) is precisely coupled to mechanical execution (Lzts1 at AJs) to remodel Rootletin fibers prior to neuronal departure from the ventricular surface.

### ZikV-NS5 hijacks Rootlet fibers to trigger early neuronal delamination

ZIKV is a flavivirus transmitted by Aedes mosquitoes that can cause neonatal microcephaly (Yuan et al., 2017) (**Fig. 5A**). Its single-stranded RNA genome encodes ten mature viral proteins (Hasan et al., 2018), among which ZIKV-NS5 interacts with centrosomal proteins such as Rootletin in NE cells, leading to an atypical, non-genetic ciliopathy (Saade et al., 2020) (**Fig. 5B**). ZIKV-NS5 also localizes to the apical membrane and induces premature apical constriction (Saade et al., 2020) (**Fig. 5B**). By perturbing these key cellular mechanisms, ZIKV-NS5 drives premature neurogenesis and depletes the progenitor pool essential for CNS growth (Saade et al., 2020).

**Figure 5.**
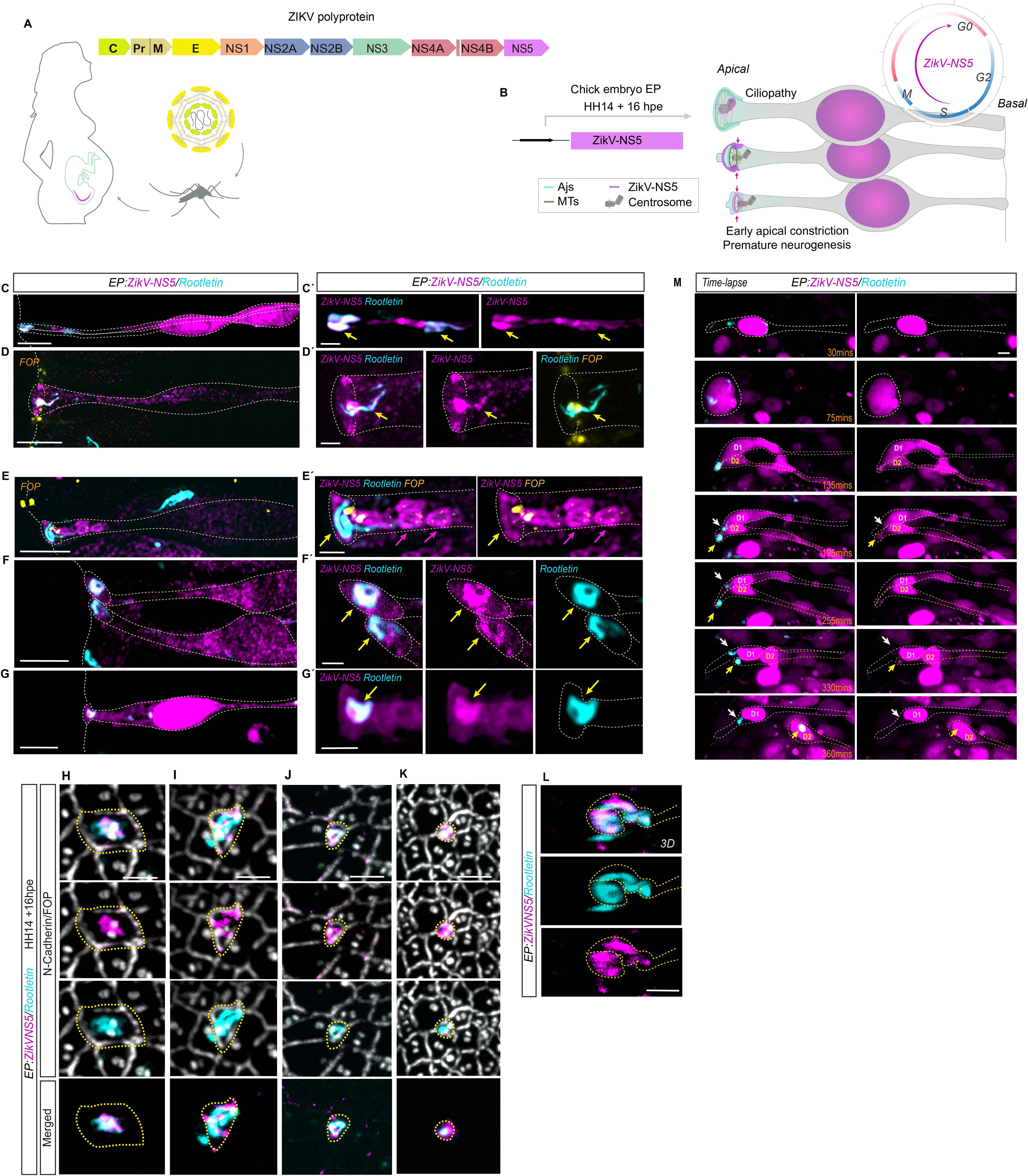
ZikV-NS5 hijacks Rootletin fibers in all their conformational startes. **(A)** Schematic of the Zika virus (ZIKV) polyprotein, showing structural (C, prM, E) and non-structural (NS1–NS5) proteins. Maternal ZIKV infection via *Aedes* mosquitoes can cause fetal brain abnormalities, including congenital microcephaly. **(B)** Schematic illustrating electroporation (EP) of the ZikV-NS5 (pCAGGS-ZikVNS5-Flag) into the developing chick NT at HH14 for fixation and immunostaining at HH17 (16hpe). The diagram summarizes NE cell behavior following ZikV-NS5 EP: ZikV-NS5 localizes to the nucleus and centrosome, induces ciliopathy leading to G0 entry, and accumulates at the apical membrane, triggering premature apical constriction and neurogenesis. **(C, D)** Representative transversal images of NE cells (outlined by a white dashed line) co-expressing ZikV-NS5-Flag together with EP-Rootletin, showing loading of the viral protein into the extended EP-Rootletin fiber at the apical feet (yellow arrows in **C**’) and at the FOP+ centrosome and ciliary rootlet (yellow arrows in **D’**). **C**’ and **D’** correspond to insets of **C** and **D**, respectively. **(E, F)** Representative transversal images of NE cells (outlined by a white dashed line) co-expressing ZikV-NS5-Flag together with EP-Rootletin, showing colocalization of ZikV-NS5-Flag with the EP-Rootletin fiber in a rim-like conformation at the apical end-surface (yellow arrows in **E’** and **F’**). Magenta arrows highlight ZikV-NS5 aggregates in **E’**. **E’** and **F’** show insets of **E** and **F**, respectively. **(G, G’)** Representative transversal images of an NE cell (outlined by a white dashed line) co-expressing ZikV-NS5-Flag together with EP-Rootletin, showing colocalization at the abscission site (yellow arrows in **G’**). **G’** insets of **G.** **(H-K)** Representative *en face* images showing apical endfeet of different surface sizes where EP ZikV-NS5-Flag and EP-Rootletin colocalize. The centrosome is stained with anti-FOP, and apical endfeet are delineated by anti-N-Cadherin. Yellow dashed lines outline the apical endfeet of highlighted cells. **(L)** 3D reconstruction of the apical endfoot shown in (H), with the yellow arrow indicating colocalization of ZikV-NS5-Flag and EP-Rootletin at the apical foot, and white arrow showing ZikV-NS5-Flag and EP-Rootletin fibers at the apical end surface. **(M)** Selected frames form an in vivo time lapse showing a transversal view of NE cell (outlined by a white dashed line) co-expressing ZikV-NS5-RFP and GFP-Rootletin. During the course of the time lapse, the cell undergoes mitosis. The white arrow points to ZikV-NS5-RFP and GFP-Rootletin at the apical endfoot of the daughter cells, while the yellow arrow marks the position of the delaminating apical endfoot of one of the daughter cells at different time points. Time is indicated in minutes (mins). (**Movie EV6**) Scale bars 5 μm (**C-G, L**), 2 μm (**Ć-G’, H-L**).

In light of our previous finding that ZIKV-NS5 interacts with Rootletin, together with our new insights into Rootletin fiber dynamics during apical constriction in NE cells, we next examined the topology of ZikV-NS5 in relation to Rootletin fiber remodeling by mis-expressing pCAGGS-ZikV-NS5-Flag in NE cells. In HH17 chick embryo NTs at 16 hpe, ZikV-NS5 displayed several subcellular distributions that overlapped with Rootletin fiber remodeling (**Fig. 5C-G**). In some NE cells, ZikV-NS5 localized at the nucleus and colocalized with Rootletin fibers extending throughout the apical process toward the apical endfoot (yellow arrows, **Fig. 5C, C’**) and with the ciliary rootlet beneath the FOP+ centrosome (yellow arrows, **Fig. 5 D, D’**). In other cells, ZikV-NS5 multimerization generated ring-like nuclear aggregates that exited the nucleus and intercalated with the extended apical Rootletin fibers (Saade et al., 2020) (purple arrows, **Fig. 5E, E’**). Additional NE cells showed ZikV-NS5 colocalizing with Rootletin fibers in rim-like conformations at the apical end surface (yellow arrows, **Fig. 5E, F**) and at the abscission site, which appeared prematurely in Zikv-NS5 expressing cells when the nucleus remained close to the ventricular surface (yellow arrow, **Fig. 5G, G’**).

To further characterize these patterns, we analyzed ZikV-NS5 and Rootletin distribution at the apical end surface by *en face* imaging, labeling centrosomes with FOP and the apical junction with N-Cadherin (**Fig. 5G-J**). In cells with relatively large apical domains, ZikV-NS5 colocalized with Rootletin at the centrosome (**Fig. 5H**). When Rootletin fibers expanded at the apical endfoot, ZikV-NS5 accumulated along these fibers (**Fig. 5I**). A 3D reconstruction of the same cell confirmed this colocalization, showing ZikV-NS5 and Rootletin fibers in a rim-like configuration at the apical endfoot and extending upwards toward the apical process (**Fig. 5L**). In constricting apical endfeet, ZikV-NS5 aligned beneath the apical junction, overlapping with the Rootletin rim (**Fig. 5J**). Upon full apical constriction, presumably immediately before abscission, ZikV-NS5 with Rootletin fibers coalesced into a single apical vertex (**Fig. 5K**).

We next used live tissue imaging to monitor the impact of ZikV-NS5 on NE cell delamination in relation to Rootletin fiber subcellular distribution (Fig. **5M**, Movie **EV6**). Following mitosis, ZikV-NS5 localised at the apical end foot of both daughter cells. The daughter cell with concentrated ZikV-NS5 and Rootletin fibers at the apical end-foot delaminated prematurely (yellow arrow) when the nucleus was still in proximity to the ventricular surface, whereas cells in which Rootletin fiber remained extended at the apical foot did not delaminate (white arrow). Together, these observations suggest that ZikV-NS5 might exploit changes in Rootletin conformation and subcellular localization to hijack key mechanisms of apical constriction and delamination, offering insight into the mechanisms by which ZikaV infection induces premature neurogenesis.

## DISCUSSION

In this study, we provide the first characterization of Rootletin localisation and dynamic behavior in neuroepithelial (NE) cells, and describe a potential mechanical role of Rootletin in NE delamination in neurogenesis. We show that Rootletin fibers are not static structures solely resisting mechanical stress, but instead display dynamic and flexible properties. In NE cells, Rootletin fibers extend beyond the centriolar linker and ciliary rootlet, spanning the entire apical foot toward the nucleus and exhibiting apical-to-basal bidirectional movement (**Fig. 1**). Because Rootletin fibers can indirectly interact with the nuclear envelope through cytoskeletal linkers such as Nesprins and SUN proteins (Potter et al., 2017), their motion along the ventricular surface may be coordinated with interkinetic nuclear migration. In mitotic cells, Rootletin localizes to spindle poles either symmetrically or asymmetrically; the asymmetric pattern likely corresponds to the mother centrosome, which retains the ciliary remnant (Paridaen et al., 2013; Saade et al., 2013) and the microtubule-anchoring protein Ninein (Wang et al., 2009), a component of the CEP68–CNAP1–Rootletin complex essential for centriole cohesion and microtubule organization (Theile et al., 2023).

We further examined the apical process of NE cells and observed that, in some cells, the Rootlet fibers retract toward the apical endfoot, extending then from the centrosome in a rim-like organization toward one side of the apical endfoot cortex (**Fig. 2**). Consistent with this observation, recent studies show that Rootlet fibers possess flexible protrusions and lateral filament networks that allow adaptive deformation and tethering to intracellular membranes (van Hoorn & Carter, 2024). Our findings further support that the Rootletin fibers undergo regulated structural remodeling, suggesting an essential role in shaping NE cell architecture.

Rootletin fibers at the apical endfoot of NE cells first remodel into a rim-like structure, followed by remodeling of the apicolateral membrane and further constriction of both structures (**Fig. 2**). This stepwise reduction of the apical endfoot occurs in coordination with key cytoskeletal changes (**Fig. 3**). First, stable microtubules at the apical endfoot assemble in an anisotropic manner at the same region where the Rootletin fiber rim is located, delineating the new apical surface. This is followed by an enrichment of Lzts1 and Drebrin at the apical endfoot cortex, along with an increase in contractile actomyosin activity. We propose that adherens junction (AJs) turnover occurs during the first step through tubulin acetylation, as its disruption in other cellular contexts blocks contractile F-actin formation at cell–cell contacts and leads to internalization of AJ proteins (Ivanov et al., 2006). In the second step, irreversible AJ disassembly is regulated by Lzts1 to promote neuronal delamination (Kawaguchi, 2020; Kawaue et al., 2019). Moreover, microtubule acetylation has been shown to slow and delay abscission during cell division, highlighting microtubules as key drivers of abscission dynamics (Kodba et al., 2025). This leads us to suggest that Rootletin fiber remodeling accompanied by tubulin acetylation ensures the precise timing of apical surface shape changes, preventing premature or delayed remodeling, maintaining epithelial integrity, and avoiding developmental defects.

In multiciliated cells, Rootletin fibers have been shown to interact with actin through the Ezrin–Fhod3 complex (Yasunaga et al., 2022). Ezrin acts as a membrane–actin linker, resisting mechanical stress and preventing blebbing during contractility, as evidenced by apical membrane destabilization in Ezrin knockout models (Korkmazhan & Dunn, 2022; Shinozuka et al., 2025). Fhod3, on the other hand, induces apical constriction by nucleating and assembling F-actin at the apical junctional complex (AJC) in NE cells, driving neural plate bending and tube closure (Sulistomo et al., 2019). Thus, Rootletin fibers may scaffold the apical actomyosin network via Ezrin–Fhod3, channeling contractile forces from Fhod3-nucleated actin to the membrane and junctions. In the neuroepithelium, this mechanism could fine-tune apical membrane remodeling and prevent excessive deformation during neurogenic apical constriction.

We further demonstrated that Rootletin fiber organization is maintained in both mouse and human neuroepithelium, highlighting its evolutionarily conserved role in NE cell morphogenesis **(Fig. 1)**. Interestingly, the human CROCC gene, which encodes Rootletin, underwent segmental duplication during evolution, giving rise to the human-specific paralog CROCCP2 (Van Heurck et al., 2023). CROCCP2 has been implicated in the expansion of basal progenitors during human corticogenesis, potentially contributing to increased brain size and complexity through links with primary cilia dynamics and mTOR signaling (Van Heurck et al., 2023). Apical constriction, however, is critical for the proper delamination of basal progenitor cells from the ventricular zone (VZ); failure in this process leads to VZ overcrowding, disrupted cortical layering, and microcephaly-like phenotypes (Kawaguchi, 2020). It will be important to investigate whether the protein encoded by CROCCP2 retains its oligomerization capacity and whether it exerts species-specific effects on Rootlet fiber remodeling. Future studies should explore how CROCC and CROCCP2 function together in the context of human neurogenesis.

In the context of Zikopathy, we have shown that ZikV-NS5 hijacks Rootletin fibers in all their conformational states—from the extended form at the apical foot to the rim-like conformation—suggesting a potential additional mechanism by which ZikV promotes premature neuron delamination (**Fig. 5**). Viruses in other systems have similarly been reported to exploit cytoskeletal remodeling to facilitate spreading and, in some cases, pathologically accelerate constriction (Lin et al., 2021; Naghavi, 2021; Stephens & Naghavi, 2024). For example, respiratory viruses induce apical infection in epithelia, activating RhoA/ROCK signaling, pulsed myosin II contractility, AJ protein endocytosis, and cell extrusion (Lin et al., 2021). Given that Rootletin fibers link the centrosome to the nuclear envelope, we propose that they may serve as an entry point for ZikV-NS5 translocation into the apical end foot. To test this hypothesis, it will be important to identify clusters of key residues mediating protein–protein interactions between ZikV-NS5 and Rootletin. By analyzing point mutations, we could potentially prevent apical translocation, thereby preserving cell fate determinants and maintaining essential mechanisms for NE cell proliferation and tissue integrity during CNS development. These experiments would provide mechanistic insight into ZIKV pathogenesis and inform the design of antiviral strategies targeting the molecular interfaces involved in ZikV-NS5–Rootletin interactions.

## ACKNOWLEDGMENTS

We are grateful to researchers that kindly provided antibodies and DNAs, Flag-Rootletin was provided from Pr. Kunsoo Rhee (Seoul National University), Rootletin-mScarlet was provided by Dr. Robert Mahen (University of Leicester). We thank D. Diego S. Ferrero (IBMB-CSIC) for generating plasmid derivatives, GFP-Rootletin and pCAGGS-ZikV-NS5-RFP. We thank Dr. Mariona Arbonés and Dr. Maria José Barallobre (IBMB-CSIC) for providing fixed mouse embryos, and for Pr. Elisa Martí for valuable feedback and discussion on the manuscript. The work in MS’s laboratory was supported by the grants CNS2023-144942 (Proyectos de Generación de Conocimiento 2022), PID2019-110157RA-I00, and RED2022-134100-T. AW and GCG are recipients of Juan de la Cierva fellowships FJC2021-047012-I and JDC2023-051684-I, respectively. PEB was supported by a FI SDUR 2021 fellowship FISDU-00400.

## AUTHOR CONTRIBUTIONS

**AW** and **GCG** developed the methodology, conceived and performed most of the experiments, analyzed the data, discussed the results, and revised the manuscript.

**PEB** generated the human 3D neural organoids, analyzed the data, discussed the results, and revised the manuscript.

**SU** provided technical assistance.

**MS** supervised the laboratory members, conceived the experiments, analyzed the data, discussed the results, and drafted the manuscript.

## References

Alvarez, Y. D., van der Spuy, M., Wang, J. X., Noordstra, I., Tan, S. Z., Carroll, M., Yap, A. S., Serralbo, O., & White, M. D. (2024). A Lifeact-EGFP quail for studying actin dynamics in vivo. J Cell Biol, 223(9). 10.1083/jcb.202404066

Au, F. K., Jia, Y., Jiang, K., Grigoriev, I., Hau, B. K., Shen, Y., Du, S., Akhmanova, A., & Qi, R. Z. (2017). GAS2L1 Is a Centriole-Associated Protein Required for Centrosome Dynamics and Disjunction. Dev Cell, 40(1), 81–94. 10.1016/j.devcel.2016.11.019

Baek, C., Freem, L., Goiame, R., Sang, H., Morin, X., & Tozer, S. (2018). Mib1 prevents Notch Cis-inhibition to defer differentiation and preserve neuroepithelial integrity during neural delamination. PLoS Biol, 16(4), e2004162. 10.1371/journal.pbio.2004162

Bocanegra-Moreno, L., Singh, A., Hannezo, E., Zagorski, M., & Kicheva, A. (2023). Cell cycle dynamics control fluidity of the developing mouse neuroepithelium. Nat Phys, 19(7), 1050–1058. 10.1038/s41567-023-01977-w

Bosveld, F., Wang, Z., & Bellaiche, Y. (2018). Tricellular junctions: a hot corner of epithelial biology. Curr Opin Cell Biol, 54, 80–88. 10.1016/j.ceb.2018.05.002

Buchman, J. J., Tseng, H. C., Zhou, Y., Frank, C. L., Xie, Z., & Tsai, L. H. (2010). Cdk5rap2 interacts with pericentrin to maintain the neural progenitor pool in the developing neocortex. Neuron, 66(3), 386–402. 10.1016/j.neuron.2010.03.036

Camargo Ortega, G., Falk, S., Johansson, P. A., Peyre, E., Broix, L., Sahu, S. K., Hirst, W., Schlichthaerle, T., De Juan Romero, C., Draganova, K., Vinopal, S., Chinnappa, K., Gavranovic, A., Karakaya, T., Steininger, T., Merl-Pham, J., Feederle, R., Shao, W., Shi, S. H., Gotz, M. (2019). The centrosome protein AKNA regulates neurogenesis via microtubule organization. Nature, 567(7746), 113–117. 10.1038/s41586-019-0962-4

Chen, J. V., Kao, L. R., Jana, S. C., Sivan-Loukianova, E., Mendonca, S., Cabrera, O. A., Singh, P., Cabernard, C., Eberl, D. F., Bettencourt-Dias, M., & Megraw, T. L. (2015). Rootletin organizes the ciliary rootlet to achieve neuron sensory function in Drosophila. J Cell Biol, 211(2), 435–453. 10.1083/jcb.201502032

Conroy, P. C., Saladino, C., Dantas, T. J., Lalor, P., Dockery, P., & Morrison, C. G. (2012). C-NAP1 and rootletin restrain DNA damage-induced centriole splitting and facilitate ciliogenesis. Cell Cycle, 11(20), 3769–3778. 10.4161/cc.21986

Das, R. M., & Storey, K. G. (2014). Apical abscission alters cell polarity and dismantles the primary cilium during neurogenesis. Science, 343(6167), 200–204. 10.1126/science.1247521

Doobin, D. J., Kemal, S., Dantas, T. J., & Vallee, R. B. (2016). Severe NDE1-mediated microcephaly results from neural progenitor cell cycle arrests at multiple specific stages. Nat Commun, 7, 12551. 10.1038/ncomms12551

Ferrero, D. S., Ruiz-Arroyo, V. M., Soler, N., Uson, I., Guarne, A., & Verdaguer, N. (2019). Supramolecular arrangement of the full-length Zika virus NS5. PLoS Pathog, 15(4), e1007656. 10.1371/journal.ppat.1007656

Francis, F., & Cappello, S. (2021). Neuronal migration and disorders - an update. Curr Opin Neurobiol, 66, 57–68. 10.1016/j.conb.2020.10.002

Frith, T. J. R., Briscoe, J., & Boezio, G. L. M. (2024). From signalling to form: the coordination of neural tube patterning. Curr Top Dev Biol, 159, 168–231. 10.1016/bs.ctdb.2023.11.004

Gilliam, J. C., Chang, J. T., Sandoval, I. M., Zhang, Y., Li, T., Pittler, S. J., Chiu, W., & Wensel, T. G. (2012). Three-dimensional architecture of the rod sensory cilium and its disruption in retinal neurodegeneration. Cell, 151(5), 1029–1041. 10.1016/j.cell.2012.10.038

Gonzalez-Gobartt, E., Blanco-Ameijeiras, J., Usieto, S., Allio, G., Benazeraf, B., & Marti, E. (2021). Cell intercalation driven by SMAD3 underlies secondary neural tube formation. Dev Cell, 56(8), 1147–1163 e1146. 10.1016/j.devcel.2021.03.023

Hasan, S. S., Sevvana, M., Kuhn, R. J., & Rossmann, M. G. (2018). Structural biology of Zika virus and other flaviviruses. Nat Struct Mol Biol, 25(1), 13–20. 10.1038/s41594-017-0010-8

Hatta, K., & Takeichi, M. (1986). Expression of N-cadherin adhesion molecules associated with early morphogenetic events in chick development. Nature, 320(6061), 447–449. 10.1038/320447a0

Heng, J. I., Nguyen, L., Castro, D. S., Zimmer, C., Wildner, H., Armant, O., Skowronska-Krawczyk, D., Bedogni, F., Matter, J. M., Hevner, R., & Guillemot, F. (2008). Neurogenin 2 controls cortical neuron migration through regulation of Rnd2. Nature, 455(7209), 114–118. 10.1038/nature07198

Herrera, A., Menendez, A., Torroba, B., Ochoa, A., & Pons, S. (2021). Dbnl and beta-catenin promote pro-N-cadherin processing to maintain apico-basal polarity. J Cell Biol, 220(6). 10.1083/jcb.202007055

Huangfu, D., & Anderson, K. V. (2005). Cilia and Hedgehog responsiveness in the mouse. Proc Natl Acad Sci U S A, 102(32), 11325–11330. 10.1073/pnas.0505328102

Ivanov, A. I., McCall, I. C., Babbin, B., Samarin, S. N., Nusrat, A., & Parkos, C. A. (2006). Microtubules regulate disassembly of epithelial apical junctions. BMC Cell Biol, 7, 12. 10.1186/1471-2121-7-12

Jimenez, A. J., Schaeffer, A., De Pascalis, C., Letort, G., Vianay, B., Bornens, M., Piel, M., Blanchoin, L., & Thery, M. (2021). Acto-myosin network geometry defines centrosome position. Curr Biol, 31(6), 1206–1220 e1205. 10.1016/j.cub.2021.01.002

Kasioulis, I., Das, R. M., & Storey, K. G. (2017). Inter-dependent apical microtubule and actin dynamics orchestrate centrosome retention and neuronal delamination. Elife, 6. 10.7554/eLife.26215

Kawaguchi, A. (2020). Neuronal Delamination and Outer Radial Glia Generation in Neocortical Development. Front Cell Dev Biol, 8, 623573. 10.3389/fcell.2020.623573

Kawaue, T., Sagou, K., Kiyonari, H., Ota, K., Okamoto, M., Shinoda, T., Kawaguchi, A., & Miyata, T. (2014). Neurogenin2-d4Venus and Gadd45g-d4Venus transgenic mice: visualizing mitotic and migratory behaviors of cells committed to the neuronal lineage in the developing mammalian brain. Dev Growth Differ, 56(4), 293–304. 10.1111/dgd.12131

Kawaue, T., Shitamukai, A., Nagasaka, A., Tsunekawa, Y., Shinoda, T., Saito, K., Terada, R., Bilgic, M., Miyata, T., Matsuzaki, F., & Kawaguchi, A. (2019). Lzts1 controls both neuronal delamination and outer radial glial-like cell generation during mammalian cerebral development. Nat Commun, 10(1), 2780. 10.1038/s41467-019-10730-y

Ko, D., Kim, J., Rhee, K., & Choi, H. J. (2020). Identification of a Structurally Dynamic Domain for Oligomer Formation in Rootletin. J Mol Biol, 432(13), 3915–3932. 10.1016/j.jmb.2020.04.012

Kodba, S., Oztop, A., van Berkum, E., Katrukha, E. A., Iwanski, M. K., Nijenhuis, W., Kapitein, L. C., & Chaigne, A. (2025). Aurora B controls microtubule stability to regulate abscission dynamics in stem cells. Cell Rep, 44(2), 115238. 10.1016/j.celrep.2025.115238

Korkmazhan, E., & Dunn, A. R. (2022). The membrane-actin linker ezrin acts as a sliding anchor. Sci Adv, 8(31), eabo2779. 10.1126/sciadv.abo2779

Le Dreau, G., Saade, M., Gutierrez-Vallejo, I., & Marti, E. (2014). The strength of SMAD1/5 activity determines the mode of stem cell division in the developing spinal cord. J Cell Biol, 204(4), 591–605. 10.1083/jcb.201307031

Lin, W. W., Tsay, A. J., Lalime, E. N., Pekosz, A., & Griffin, D. E. (2021). Primary differentiated respiratory epithelial cells respond to apical measles virus infection by shedding multinucleated giant cells. Proc Natl Acad Sci U S A, 118(11). 10.1073/pnas.2013264118

Long, K., Moss, L., Laursen, L., Boulter, L., & Ffrench-Constant, C. (2016). Integrin signalling regulates the expansion of neuroepithelial progenitors and neurogenesis via Wnt7a and Decorin. Nat Commun, 7, 10354. 10.1038/ncomms10354

Mahen, R. (2021). The structure and function of centriolar rootlets. J Cell Sci, 134(16). 10.1242/jcs.258544

Marthiens, V., & ffrench-Constant, C. (2009). Adherens junction domains are split by asymmetric division of embryonic neural stem cells. EMBO Rep, 10(5), 515–520. 10.1038/embor.2009.36

Martin, A. C., & Goldstein, B. (2014). Apical constriction: themes and variations on a cellular mechanism driving morphogenesis. Development, 141(10), 1987–1998. 10.1242/dev.102228

Miyata, T., Okamoto, M., Shinoda, T., & Kawaguchi, A. (2014). Interkinetic nuclear migration generates and opposes ventricular-zone crowding: insight into tissue mechanics. Front Cell Neurosci, 8, 473. 10.3389/fncel.2014.00473

Naghavi, M. H. (2021). HIV-1 capsid exploitation of the host microtubule cytoskeleton during early infection. Retrovirology, 18(1), 19. 10.1186/s12977-021-00563-3

Pacary, E., Heng, J., Azzarelli, R., Riou, P., Castro, D., Lebel-Potter, M., Parras, C., Bell, D. M., Ridley, A. J., Parsons, M., & Guillemot, F. (2011). Proneural transcription factors regulate different steps of cortical neuron migration through Rnd-mediated inhibition of RhoA signaling. Neuron, 69(6), 1069–1084. 10.1016/j.neuron.2011.02.018

Pacary, E., Martynoga, B., & Guillemot, F. (2012). Crucial first steps: the transcriptional control of neuron delamination. Neuron, 74(2), 209–211. 10.1016/j.neuron.2012.04.002

Paridaen, J. T., Wilsch-Brauninger, M., & Huttner, W. B. (2013). Asymmetric inheritance of centrosome-associated primary cilium membrane directs ciliogenesis after cell division. Cell, 155(2), 333–344. 10.1016/j.cell.2013.08.060

Park, H. C., Kim, C. H., Bae, Y. K., Yeo, S. Y., Kim, S. H., Hong, S. K., Shin, J., Yoo, K. W., Hibi, M., Hirano, T., et al. (2000). Analysis of upstream elements in the HuC promoter leads to the establishment of transgenic zebrafish with fluorescent neurons. Dev Biol 227, 279–293. 10.1006/dbio.2000.9898

Potter, C., Zhu, W., Razafsky, D., Ruzycki, P., Kolesnikov, A. V., Doggett, T., Kefalov, V. J., Betleja, E., Mahjoub, M. R., & Hodzic, D. (2017). Multiple Isoforms of Nesprin1 Are Integral Components of Ciliary Rootlets. Curr Biol, 27(13), 2014–2022 e2016. 10.1016/j.cub.2017.05.066

Ranie, S. N., & White, M. D. (2025). Apical constriction in morphogenesis: From actomyosin architecture to regulatory networks. Curr Opin Cell Biol, 95, 102562. 10.1016/j.ceb.2025.102562

Rehm, K., Panzer, L., van Vliet, V., Genot, E., & Linder, S. (2013). Drebrin preserves endothelial integrity by stabilizing nectin at adherens junctions. J Cell Sci, 126(Pt 16), 3756–3769. 10.1242/jcs.129437

Rousso, D. L., Pearson, C. A., Gaber, Z. B., Miquelajauregui, A., Li, S., Portera-Cailliau, C., Morrisey, E. E., & Novitch, B. G. (2012). Foxp-mediated suppression of N-cadherin regulates neuroepithelial character and progenitor maintenance in the CNS. Neuron, 74(2), 314–330. 10.1016/j.neuron.2012.02.024

Saade, M., Ferrero, D. S., Blanco-Ameijeiras, J., Gonzalez-Gobartt, E., Flores-Mendez, M., Ruiz-Arroyo, V. M., Martinez-Saez, E., Ramon, Y. C. S., Akizu, N., Verdaguer, N., & Marti, E. (2020). Multimerization of Zika Virus-NS5 Causes Ciliopathy and Forces Premature Neurogenesis. Cell Stem Cell, 27(6), 920–936 e928. 10.1016/j.stem.2020.10.002

Saade, M., Gonzalez-Gobartt, E., Escalona, R., Usieto, S., & Marti, E. (2017). Shh-mediated centrosomal recruitment of PKA promotes symmetric proliferative neuroepithelial cell division. Nat Cell Biol, 19(5), 493–503. 10.1038/ncb3512

Saade, M., Gutierrez-Vallejo, I., Le Dreau, G., Rabadan, M. A., Miguez, D. G., Buceta, J., & Marti, E. (2013). Sonic hedgehog signaling switches the mode of division in the developing nervous system. Cell Rep, 4(3), 492–503. 10.1016/j.celrep.2013.06.038

Saade, M., & Marti, E. (2025). Early spinal cord development: from neural tube formation to neurogenesis. Nat Rev Neurosci, 26(4), 195–213. 10.1038/s41583-025-00906-5

Shinozuka, T., Okubo, T., Sasai, N., & Takada, S. (2025). Wnt-dependent mechanism of the apical constriction of roof plate cells in developing mouse spinal cord. Front Cell Dev Biol, 13, 1571770. 10.3389/fcell.2025.1571770

Stephens, C., & Naghavi, M. H. (2024). The host cytoskeleton: a key regulator of early HIV-1 infection. FEBS J, 291(9), 1835–1848. 10.1111/febs.16706

Styczynska-Soczka, K., & Jarman, A. P. (2015). The Drosophila homologue of Rootletin is required for mechanosensory function and ciliary rootlet formation in chordotonal sensory neurons. Cilia, 4, 9. 10.1186/s13630-015-0018-9

Sulistomo, H. W., Nemoto, T., Yanagita, T., & Takeya, R. (2019). Formin homology 2 domain-containing 3 (Fhod3) controls neural plate morphogenesis in mouse cranial neurulation by regulating multidirectional apical constriction. J Biol Chem, 294(8), 2924–2934. 10.1074/jbc.RA118.005471

Theile, L., Li, X., Dang, H., Mersch, D., Anders, S., & Schiebel, E. (2023). Centrosome linker diversity and its function in centrosome clustering and mitotic spindle formation. EMBO J, 42(17), e109738. 10.15252/embj.2021109738

Turkyilmaz, A., & Sager, S. G. (2022). Two New Cases of Primary Microcephaly with Neuronal Migration Defect Caused by Truncating Mutations in the ASPM Gene. Mol Syndromol, 13(1), 56–63. 10.1159/000516201

Van Heurck, R., Bonnefont, J., Wojno, M., Suzuki, I. K., Velez-Bravo, F. D., Erkol, E., Nguyen, D. T., Herpoel, A., Bilheu, A., Beckers, S., Ledent, C., & Vanderhaeghen, P. (2023). CROCCP2 acts as a human-specific modifier of cilia dynamics and mTOR signaling to promote expansion of cortical progenitors. Neuron, 111(1), 65–80 e66. 10.1016/j.neuron.2022.10.018

van Hoorn, C., & Carter, A. P. (2024). A cryo-electron tomography study of ciliary rootlet organization. Elife, 12. 10.7554/eLife.91642

Vlijm, R., Li, X., Panic, M., Ruthnick, D., Hata, S., Herrmannsdorfer, F., Kuner, T., Heilemann, M., Engelhardt, J., Hell, S. W., & Schiebel, E. (2018). STED nanoscopy of the centrosome linker reveals a CEP68-organized, periodic rootletin network anchored to a C-Nap1 ring at centrioles. Proc Natl Acad Sci U S A, 115(10), E2246–E2253. 10.1073/pnas.1716840115

Wang, X., Tsai, J. W., Imai, J. H., Lian, W. N., Vallee, R. B., & Shi, S. H. (2009). Asymmetric centrosome inheritance maintains neural progenitors in the neocortex. Nature, 461(7266), 947–955. 10.1038/nature08435

Wilmerding, A., Espana-Bonilla, P., Giakoumakis, N. N., & Saade, M. (2023). Expansion microscopy of the chick embryo neural tube to overcome molecular crowding at the centrosomes-cilia. STAR Protoc, 4(1), 101997. 10.1016/j.xpro.2022.101997

Wilmerding, A., Rinaldi, L., Caruso, N., Lo Re, L., Bonzom, E., Saurin, A. J., Graba, Y., & Delfini, M. C. (2021). HoxB genes regulate neuronal delamination in the trunk neural tube by controlling the expression of Lzts1. Development, 148(4). 10.1242/dev.195404

Wilsch-Brauninger, M., Peters, J., Paridaen, J. T., & Huttner, W. B. (2012). Basolateral rather than apical primary cilia on neuroepithelial cells committed to delamination. Development, 139(1), 95–105. 10.1242/dev.069294

Yang, J., Gao, J., Adamian, M., Wen, X. H., Pawlyk, B., Zhang, L., Sanderson, M. J., Zuo, J., Makino, C. L., & Li, T. (2005). The ciliary rootlet maintains long-term stability of sensory cilia. Mol Cell Biol, 25(10), 4129–4137. 10.1128/MCB.25.10.4129-4137.2005

Yang, J., Liu, X., Yue, G., Adamian, M., Bulgakov, O., & Li, T. (2002). Rootletin, a novel coiled-coil protein, is a structural component of the ciliary rootlet. J Cell Biol, 159(3), 431–440. 10.1083/jcb.200207153

Yasunaga, T., Wiegel, J., Bergen, M. D., Helmstadter, M., Epting, D., Paolini, A., Cicek, O., Radziwill, G., Engel, C., Brox, T., Ronneberger, O., Walentek, P., Ulbrich, M. H., & Walz, G. (2022). Microridge-like structures anchor motile cilia. Nat Commun, 13(1), 2056. 10.1038/s41467-022-29741-3

Yuan, L., Huang, X. Y., Liu, Z. Y., Zhang, F., Zhu, X. L., Yu, J. Y., Ji, X., Xu, Y. P., Li, G., Li, C., Wang, H. J., Deng, Y. Q., Wu, M., Cheng, M. L., Ye, Q., Xie, D. Y., Li, X. F., Wang, X., Shi, W., Qin, C. F. (2017). A single mutation in the prM protein of Zika virus contributes to fetal microcephaly. Science, 358(6365), 933–936. 10.1126/science.aam7120

