## Supplementary Figures and Legends for "Rootletin Fiber Dynamics Integrate Cytoskeletal Programs to Shape Neuroepithelial Architecture"

**Supplementary information:**

- **Expanded view Figures EV1 and EV3**
- **Expanded view Figure and Movie legends (EV1-EV6)**

**EXPANDED VIEW FIGURE LEGENDS**

**Figure EV1: Rootletin organizes fibers along the apical foot, adopting a distribution similar to the cis-Golgi.**

- **(A)** Transverse chick embryo NT sections show anti-GM130 labeling the cis-Golgi and DAPI labeling nuclei in the VZ and mantle zone (MZ). Dashed lines indicate the VZ and MZ.
- **(B)** Images show EP-Rootletin forming fibers (red arrows) and anti-GM130 labeling the cis-Golgi (cyan arrows) at the apical process of NE cells in the VZ.
- **(C)** Quantification of the length of EP-Rootletin fibers and GM130+ cis-Golgi at NE cell apical processes in the VZ (n = 53 across 4 embryos; median ± S.D.).

ns: not significant. Mann-Whitney *U* test, **(C)**. Scale bar 5 µm (**A, B**).

**Movie EV2: Extended Rootlet fibers move along the apico-basal axis of the neuroepithelium.** Time-lapse 3D imaging of electroporated NE cells in the VZ shows EP-Rootletin localization at the apical endfeet, with Rootlet fibers extending along the apico-basal axis and dynamically moving in both anterograde and retrograde directions (white arrows) in coordination with interkinetic nuclear migration. Scale bar 5 µm.

**Figure EV3.- Endogenous Rootletin localizes to the centriolar linker, extends into the VZ, and associates with the apical membrane in chick NE cells.**

- **(A)** Transverse sections of chick embryo NT immunostained with anti-Rootletin, anti-Centrin, and nuclei stained with DAPI. immunostaining, Images show Rootletin localisation between the two centrioles (yellow arrows) lining the apical end surface of the VZ.
- **(B)** Fluorescence intensity (AU) profile plots of anti-Centrin and anti-Rootletin across selected centrosomes to assess the distance between Rootletin and the two centrioles.
- **(C)** Transverse sections of chick embryo NT showing anti-Rootletin localization at the VZ (yellow arrows) and electroporated Arl13b-RFP labeling primary cilia along NT lumen (outlined by a dashed line)
- **(D)** Representative image showing the transversal view of an NE cell apical process electroporated (EP)-Rootletin, immunostained Rootletin, anti-ZO-1 labeling TJs, and anti-Centrin labeling centrioles. Yellow dashed line represents the cell outline; gray dashed line outlines the NT lumen.
- **(E)** Representative images of the *en face* view of NE cells immunostained with anti-Rootletin. Grey dashed line outlines the NT lumen; yellow dashed line outlines the apical endfeet of highlighted cells.
- **(F)** *En face* images showing anti-ZO-1 labeled AJs, anti-Centrin-labeled centrioles and the subcellular localization of immunostained Rootletin. (F left) Magnified view of the region highlighted by the yellow dashed line in the right. Yellow dash line outlines the highlighted apical endfoot.
- **(G)** Fluorescence intensity (AU) profile plots of anti-ZO-1 and anti-Rootletin across the apical endfeet of selected cells to assess its localization relative to AJs at the apical membrane
- **(H)** Representative images show the *en face* view of (EP)-Rootletin and anti-Rootletin immunostaining at the apical foot of NE cells. Dashed line outlines the highlighted cell.

Scale bars 5 µm (**A, C, E**), 2 µm (**D, H, I**), 1 µm (**F**)

**Movie EV4: Rootletin fibers show different configurations at the apical endfeet of NE cells.** *En face* high spatiotemporal resolution imaging of NE cell apical endfeet, expressing membrane-targeted FP and Rootletin (cyan). In some, Rootletin localization is restricted to the center of the apical area (white arrow). In others, Rootletin emerges fibers in close vicinity of the apical membrane (green arrow); and in yet others, a clear rim-like fiber occupying the apical area (yellow arrow). Scale bar 5 µm

**Movie EV5: Live imaging of Rootletin fiber retraction, Rootletin apical rim remodeling, and apical endfoot constriction in individual neuroepithelial cells.** Live imaging was performed on chick embryos at stage HH17 (16hpe), electroporated with mb-RFP and GFP-Rootletin (cyan). A representative NE cell is shown in transverse and *en face* views., The yellow arrowhead indicates the position of the most proximal edge of the Rootletin fiber at different time points. Time is indicated in minutes (mins). Scale bars 2 µm

**Movie EV6: Live imaging of the ZikV-NS5-Rootletin interaction in a delaminating NE cell.** Live imaging was performed on chick embryos at stage HH17 (16hpe), electroporated with ZikV-NS5 (purple) and Rootletin (cyan). NE cells are and the ventricular surface are outlined by a white dashed line and are shown in transverse view. The yellow arrow indicates ZikV-NS5 and Rootletin at the apical end-foot, while the white arrow marks the position of the delaminating apical foot at different time points. Time is indicated in minutes (mins).

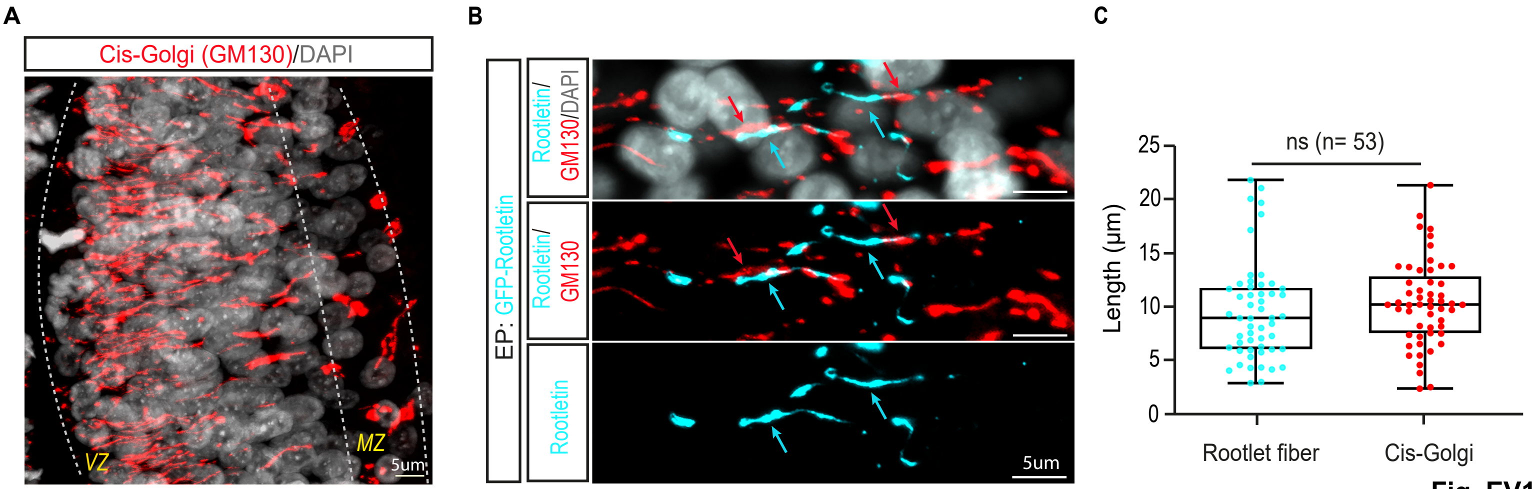

A
Cis-Golgi (GM130)/DAPI
VZ
MZ
5um
B
EP: GFP-Rootletin
Rootletin/GM130/DAPI
Rootletin/GM130
Rootletin
5um
C
ns (n= 53)
Length (μm)
25
20
15
10
5
0
Rootlet fiber
Cis-Golgi


-Fig. EV1-

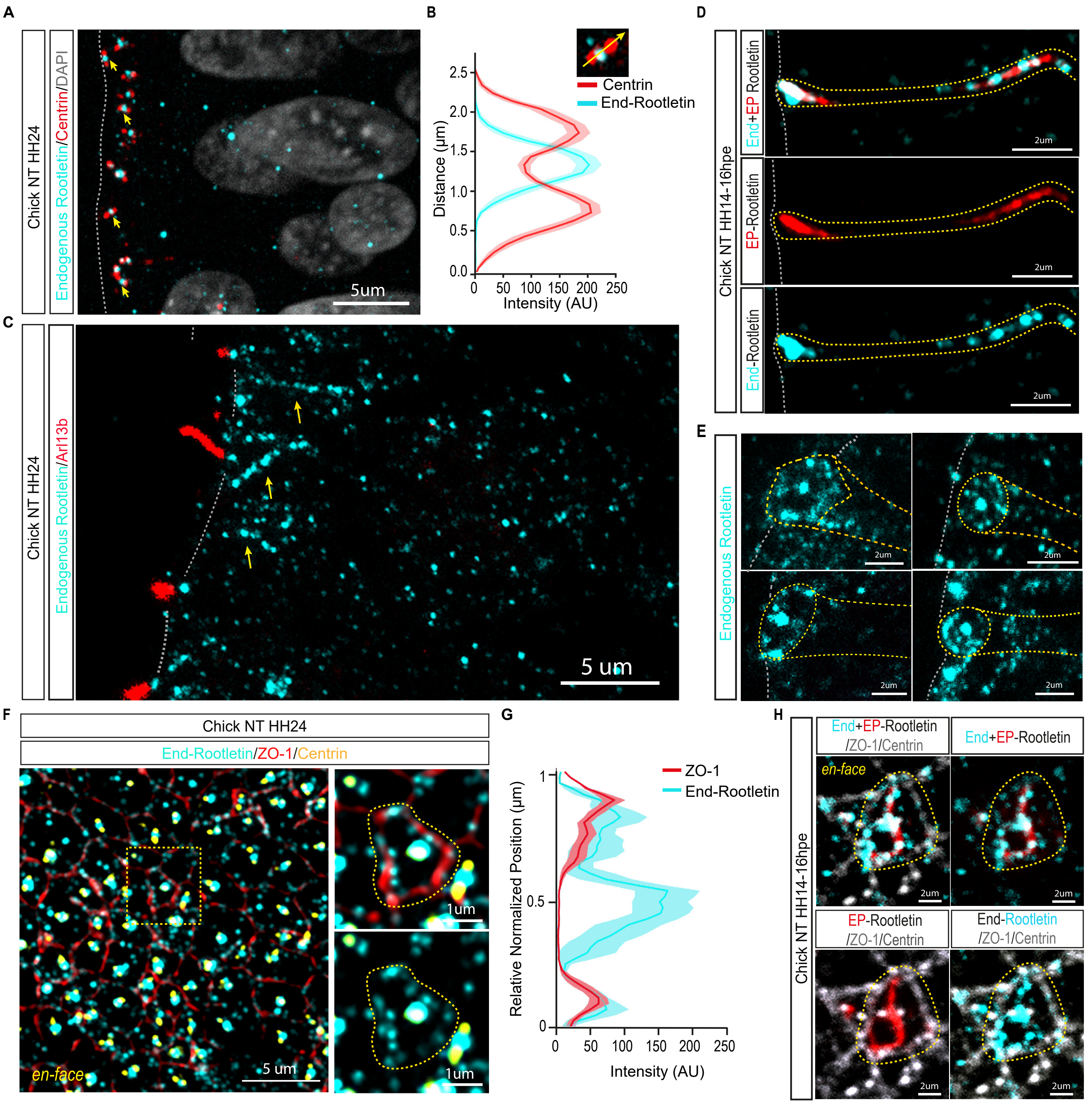

A
Chick NT HH24
Endogenous Rootletin/Centrin/DAPI
5um
B
Centrin
End-Rootletin
Distance (µm)
Intensity (AU)
D
Chick NT HH14-16hpe
End+EP Rootletin
EP-Rootletin
End-Rootletin
2um
C
Chick NT HH24
Endogenous Rootletin/Arl13b
5 um
E
Endogenous Rootletin
2um
F
Chick NT HH24
End-Rootletin/ZO-1/Centrin
en-face
5 um
1um
G
ZO-1
End-Rootletin
Relative Normalized Position (µm)
Intensity (AU)
H
Chick NT HH14-16hpe
End+EP-Rootletin /ZO-1/Centrin
End+EP-Rootletin
EP-Rootletin /ZO-1/Centrin
End-Rootletin /ZO-1/Centrin
en-face
2um


-Fig. EV3-
